# An Integrated Analysis of GLP-1R Agonist Mechanisms: Addressing Study Variations in Heterogeneous Cell Systems

**DOI:** 10.64898/2026.03.25.714139

**Authors:** Oscar Silfvergren, Sophie Rigal, Katharina Schimek, Christian Simonsson, Kajsa P Kanebratt, Felix Forschler, Burcak Yesildag, Uwe Marx, Liisa Vilén, Peter Gennemark, Gunnar Cedersund

## Abstract

Experimental cell systems support the development of pharmacological therapies such as glucagon-like peptide-1 receptor agonists (GLP-1RAs). However, their utility in drug discovery is limited due to study variability, which complicates formation of unified conclusions based on all available data. To address this, we conducted a comprehensive analysis of the GLP-1RA exenatide, incorporating 16 new and five pre-existing mono- or co-culture studies of human liver and pancreatic models. We employed a new pragmatic model-based approach designed to handle the common situation of heterogeneous *in vitro* datasets with few replicates per condition. All studies are jointly explained (disagreement<χ²-limit; 542<732), thereby providing a unified conclusion based on all studies. This work links *in vitro* biology to clinically relevant mechanisms, such as exenatide’s effect on glucose-insulin interplay, and predicts previously undescribed inter-study variabilities. Independent validation confirms predictive performance (64<83). Our new integrative approach enhances the utility of experimental cell systems in preclinical drug discovery.

## INTRODUCTION

The prevalence of type 2 diabetes mellitus (T2DM) has increased over recent decades, placing a growing burden on healthcare systems worldwide. Pharmacological therapies such as glucagon-like peptide-1 receptor agonists (GLP-1RAs) provide effective options for preventing T2DM (1,2). To support development of such drugs, experimental human cell systems can be used to study metabolic regulation and drug responses outside the human body, using e.g., conventional static cell assays and microphysiological systems (MPS). Conventional static assays are inexpensive, rapid, and well suited for generating dose-responses, but they do not capture crosstalk between cell types from different organs which limits their ability to represent pharmacological effects. In contrast, MPS are more advanced *in vitro* experimental systems, where e.g different miniaturized human organ models can be interconnected, so that such crosstalk can be emulated. Both static assays and MPS face a shared challenge: interpreting biological data is complicated by significant experimental variability arising from differences in experimental conditions such as media composition, cell types, drug exposure profiles, and donor-specific factors. Additional complicating factors include how different cell models in the same culture respond differently to identical media formulations (3), and that there generally are few donors or replicates available per condition. Importantly, it is likely that these differences are dominated by biology rather than experimental noise, since we and others have seen an excellent reproducibility across experimental cell studies (4–10). It is critical to separate true biological and mechanistic effects from observed variability to bridge *in vitro* biology and clinical responses, thereby improving the utility of experimental cell systems in drug discovery. To address this general problem in the case of T2DM, we conducted a comprehensive study of the GLP-1RA exenatide, incorporating 16 new static cell assays and MPS studies, and analysed the data jointly, using a new approach to computational modelling.

A computational model is a mathematical representation of a system and such models are increasingly used to describe dynamic biological processes (11–13). One prominent type of models is ordinary differential equation (ODE) models, which have shown potential for interpreting human cell data and drug effects (14–18). We have previously developed an ODE-based mathematical model describing human glucose–insulin interplay from clinical data (19). This work describes human-to-human variability by allowing covariables, representing for example insulin resistance and insulin production capacity, to vary freely between populations. An important alternative is to assume that covariables follow Gaussian or log-normal distributions, as is common in traditional population-based modelling (20,21). This works well when populations are large enough to estimate and validate the assumed distributions. However, in experimental cell systems, sample sizes per condition are typically small and multiple sources of variability can influence the same covariable, making such assumptions less reliable. Therefore, we analyse our GLP-1RA exenatide dataset using a new pragmatic approach tailored to heterogeneous *in vitro* datasets.

Herein, we present a comprehensive study of the GLP-1RA exenatide, incorporating 16 new mono- or co-culture studies of liver–pancreas, and analyse them using a novel model-based approach. This analysis provides a unified description of the biology across all studies by accounting for study-specific differences (**Fig. 1**). By jointly interpreting these 16 studies together with five pre-existing studies, we characterize the observed *in vitro* biology, relate the findings to clinically relevant metabolic mechanisms, and quantitatively predict previously undescribed sources of inter-study variability. This work equips the multidisciplinary scientific community with new tools to test unified hypotheses across heterogeneous datasets and advances biological understanding of *in vitro* liver-pancreas biology, thereby enhancing the utility of experimental cell systems in preclinical drug discovery.

**Figure 1:**
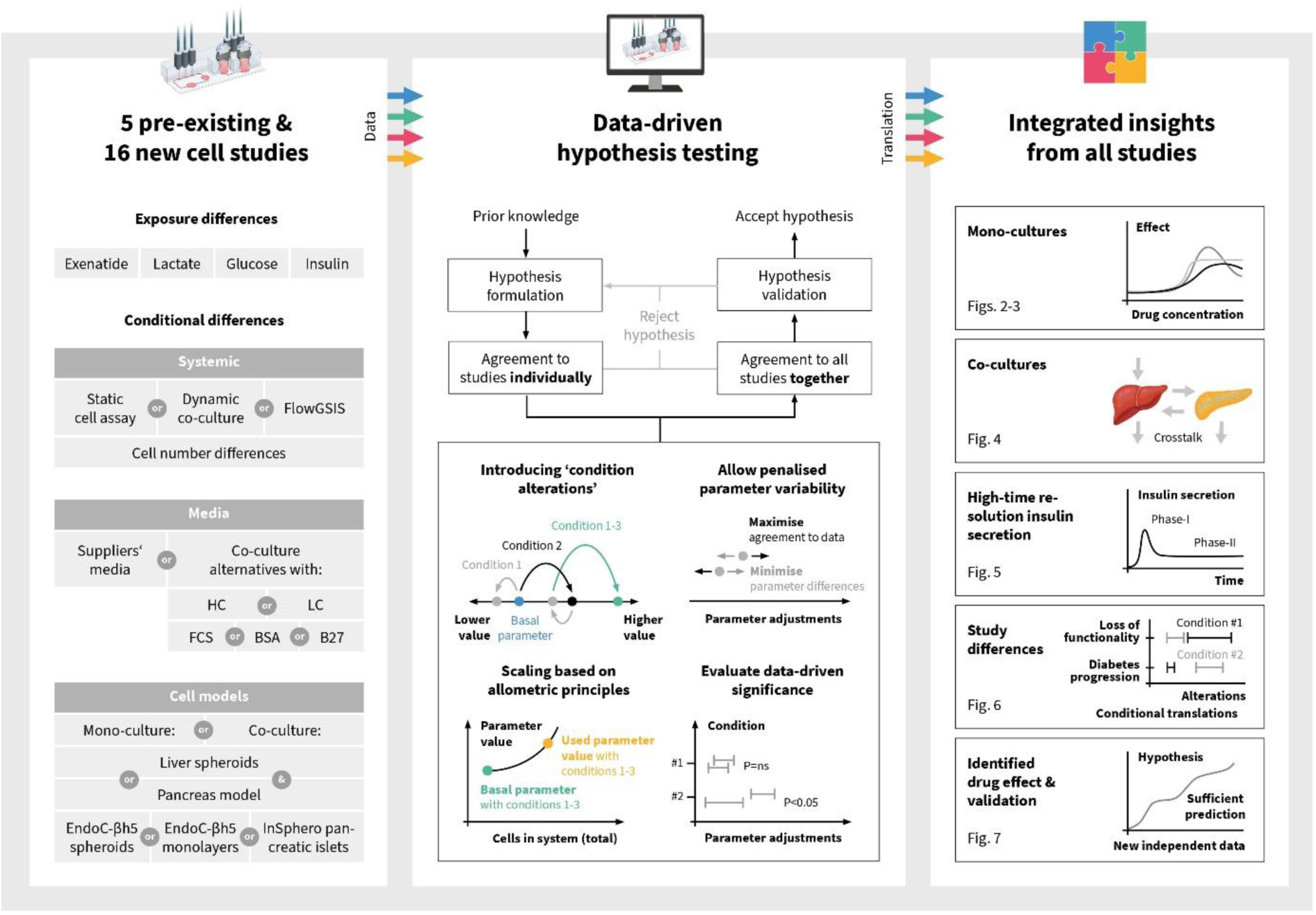
Interpreting biology and study variability for the GLP-1RA exenatide using experimental cell systems The figure illustrates study differences (left), a data-driven hypothesis testing approach (middle), and the integrated insights from all studies organized into five sections (right). Media differences include multiple formulations in which a base medium is supplemented with high or low hydrocortisone (HC or LC) and with fetal calf serum (FCS), bovine serum albumin (BSA), or B27 supplements.

## RESULTS

We conducted 16 cell-culture studies with and without the GLP-1RA exenatide (**Fig. 1, left**). The newly generated studies (**Supplementary Data S1**) were supplemented with previously reported data by us (4) and additional pre-existing studies (**Supplementary Data S2**) (22–24).

Collectively, all studies provide comprehensive insights into the effects of exenatide on *in vitro* liver-pancreas biology, as they span over a wide range of experimental conditions: differences in administered exposures, experimental system differences, media compositions, and cell models (**Fig. 1, left**). Administered exposures differed in the concentrations of exenatide (0–10^3^ nM), lactate (0–2 mM), glucose (0–20 mM), and insulin (0–100 nM). Experimental system differences comprised variations in total cell numbers (approximately 2,000 to 10⁶ cells) and the use of different experimental platforms, including static well plates, MPS (4,8–10), and a microfluidic hanging-drop-based islet perifusion system (FlowGSIS) (23,24). Media composition differences included total media volume (50–600 μL) and the use of either supplier-provided media (SM) or various co-culture media formulations. The co-culture media formulations consisted of a basal medium supplemented with either high or low hydrocortisone (HC or LC) and fetal calf serum (FCS), bovine serum albumin (BSA), or B27 supplements. Finally, cell-model differences encompassed mono- or co-cultures of liver spheroids (4,8–10) (batches 1–6), as well as multiple pancreatic models: EndoC-βH5 3D cultures (batches 1-4; spheroids), EndoC-βH5 2D monolayers (batch 5), and 3D human primary pancreatic microtissues (donors 1-11, islets) (**Fig. 1**, left).

All studies were analysed using a model-based approach (**Fig. 1**, middle). This approach tests a minimal set of hypothesised mechanisms capable of explaining all studies jointly. The key hypothesis describes how pancreatic insulin secretion is stimulated by glucose and exenatide, and how glucose utilisation increases with insulin concentration. Differences between studies are captured through mechanistic modelling that accounts for administered exposures, scaling functions reflecting differences in total cell numbers, and condition-specific parameter adjustments (condition alterations) representing biological variation across cell models, cell donors, cell-line batches, as well as their responses to different media. In brief, these condition alterations capture variation in time-dependent loss of metabolic rates (media, liver, and pancreas), insulin resistance (media and liver), and exenatide potency and efficacy (media and pancreas). The biology is encoded by 42 equations (including 10 ODEs), 44 parameters describing pancreas-hepatic metabolism, and 129 condition alterations. Parameters and condition alterations were estimated by optimizing an objective function designed to fit all studies, while keeping variability between the condition alterations low.

Using this approach, the mathematical model successfully described all cell-culture studies simultaneously (**Fig. 1**, right), as agreement with the data was not rejected by either visual assessment or a 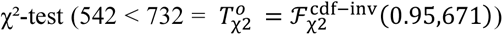. The result is presented in five sections: insulin secretion of pancreatic monocultures, glucose utilisation of hepatic monocultures, glucose-insulin interactions in pancreatic-hepatic co-cultures, sources of inter-study variability, and validation of described biology through prediction of new, independent data.

### Insulin secretion of pancreatic monocultures

We conducted static cell assays to characterize exenatide-dependent insulin secretion, glucose-dependent insulin secretion, and insulin secretion changes in media variations (**Fig. 2**).

**Figure 2:**
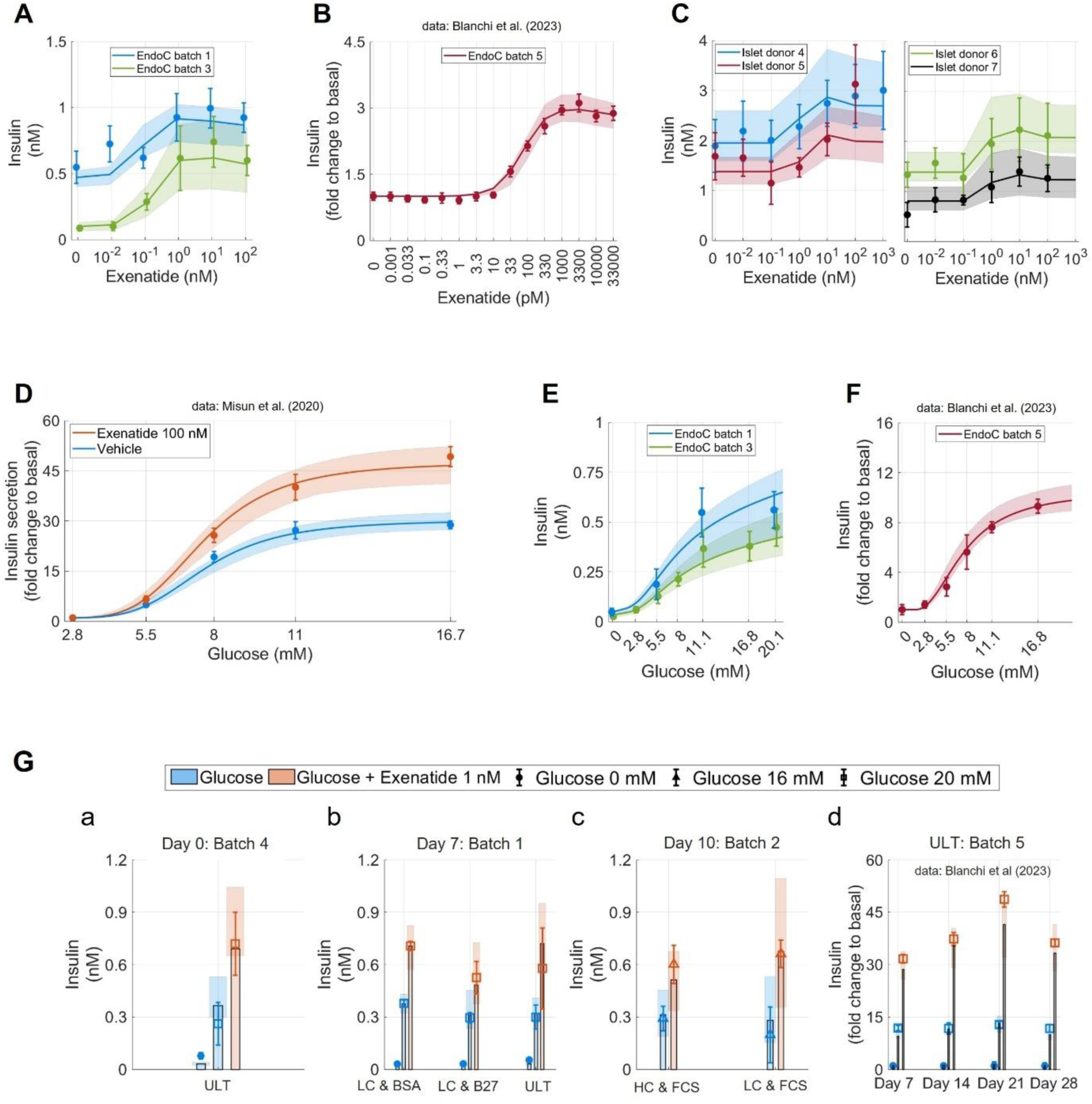
**Insulin secretion of pancreatic monocultures** Lines represent mathematical model best joint-fit to all cell-culture studies (Figs. 2-5, Supplementary Fig. S1). Shaded areas indicate model uncertainty. Error-bars represent data (mean ± SD). *(A)* EndoC-βH5 spheroid exenatide dose-response curves (Study 10: blue; Study 8: green). *(B)* EndoC-βH5 monolayer exenatide dose-response curves (Study 19) (22). *(C)* Human islet exenatide dose-response curves (Study 13). *(D)* Human islet glucose dose-response curves (Study 17) (24). Y-axis is normalised to 2.8 mM glucose group without exenatide (blue). *(E)* EndoC-βH5 spheroid glucose dose-response curves (Study 10: blue; Study 8: green). *(F)* EndoC-βH5 monolayer glucose dose-response curves (Study 19) (22). Y-axis is normalised to 0 mM glucose group. *(G)* EndoC-βH5 assays from various studies. Orange colour represents 1 nM exenatide and blue colour represents no exenatide. Marker-type show glucose concentration: 0 mM (circle), 11.1 mM (triangle), and 20 mM (square). Studies were conducted with supplier’s medium (ULT) or a basal medium supplemented with high or low hydrocortisone (HC or LC) and fetal calf serum (FCS), bovine serum albumin (BSA), or B27 supplements. a) Study 12 (spheroids). b) Day 7 of Study 7 (spheroids). c) Day 10 of Study 7 (spheroids). d) Reported data (17) of EndoC-βH5 monolayers (Study 19). Y-axis is normalised to 0 mM glucose group.

First, we found that exenatide exposure increased insulin secretion up to maximum effect (EC_100_) arguably followed by a decreasing effect if exenatide exposure was further increased, i.e., a bell-shaped dose-response (**Fig. 2A**, Endo-βH5 spheroids; **Fig. 2B**, Endo-βH5 monolayers; **Fig. 2C**, human islets). The clinical dose-response of GLP-1RAs has been reported as both sigmoidal (25) and bell-shaped in humans (26) and rats (27), indicating that a modest decline in response beyond EC₁₀₀ is plausible. An exenatide effect was observed in human islets exposed to high glucose concentrations (**Fig. 2D**; glucose above 5.5 mM), but not at low glucose concentrations (**Fig. 2D**; glucose below 5.5 mM) (24). Therefore, the effect of exenatide is implemented as acting solely on glucose-dependent insulin secretion in the mathematical model (**Supplementary Note S1**), consistent with clinical observations (28).

Second, data indicate that glucose exposure increased insulin secretion toward a maximal effect, i.e., a sigmoidal dose-response (**Fig. 2D**, Human islets; **Fig. 2E**, Endo-βH5 spheroids; **Fig. 2F**, Endo-βH5 monolayers). To adequately describe the data, a minimum threshold of 1.7 mM for stimulated glucose-dependent insulin secretion was identified (uncertainty range: 1.3-2.1 mM). This threshold was evident in pre-existing data (**Fig. 2F**) (22) and reproduced in our new studies (**Fig. 3E**). The presence of such a threshold is consistent with clinical observations, where insulin secretion approaches basal levels at approximately 3 mM glucose (29).

**Figure 3:**
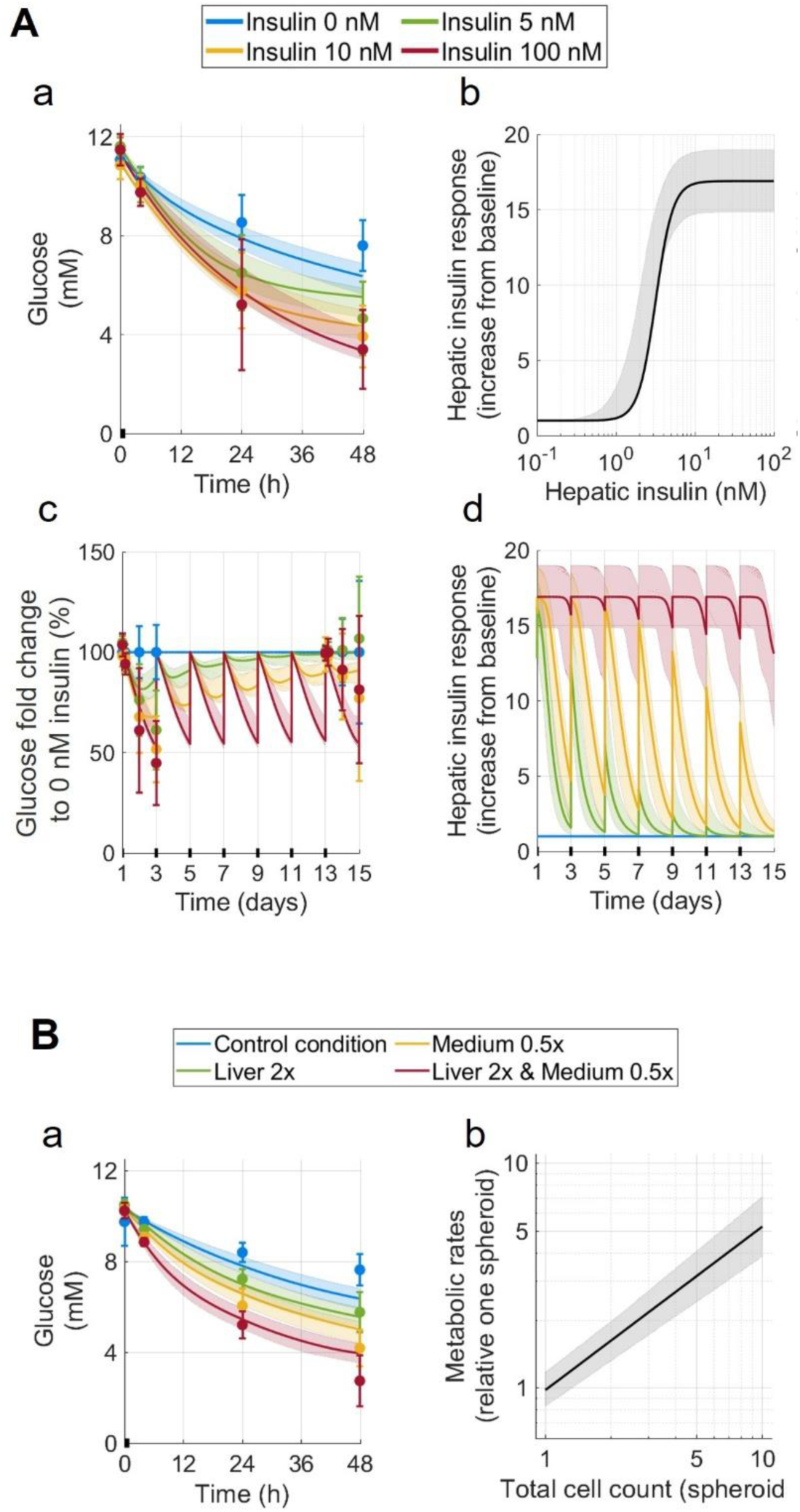
**Glucose utilisation of hepatic monocultures** Lines represent mathematical model best joint-fit to all cell-culture studies (Figs. 2-5, Supplementary Fig. S1). Shaded areas indicate model uncertainty. Error-bars represent data (mean ± SD). *(A)* Liver spheroids insulin dose response test (Study 6). a) Glucose medium concentrations days 1-3. b) Predicted insulin-dependent glucose utilisation days 1-3. c) Glucose medium fold change to non-insulin group. d) Predicted hepatic insulin-stimulated glucose utilisation throughout the full study. *(B)* Static mono-culture liver spheroid assays (Study 6). The liver spheroids are separated into four groups: Control condition (blue) denotes 40 liver spheroids in 605 µL of culture medium; 2x Liver condition (green) denotes double liver spheroids relative to control condition; 0.5x Medium condition (yellow) denotes half medium volume relative to control condition; 2x Liver & 0.5x Medium condition (red) denotes double number of liver spheroids and half medium volume relative to control condition. a) Glucose in medium. b) Metabolic rate scaling to number of spheroids.

Finally, changes in insulin secretion from EndoC-βH5 spheroids were evaluated under different media conditions over a 10-day period (**Figs. 2G a-c**). Pre-existing data from EndoC-βH5 monolayers were included in the model evaluation, which demonstrated stable insulin secretion capacity up to 28-days when cultured in the supplier’s ULT-medium (**Fig. 2G d**) (22).

Overall, the mathematical model was able to adequately represent all data on insulin secretion of pancreatic monocultures.

### Glucose utilisation of hepatic monocultures

Hepatic monocultures were used to characterize insulin-dependent glucose utilisation, insulin-independent glucose utilisation, changes in insulin sensitivity, and the dependence of glucose utilisation on the number of cells in the culture (**Fig. 3**).

First, we found that insulin exposure increased glucose utilisation (**Fig. 3A a**) with a sigmoidal dose-response (**Fig. 3A b**). The EC₅₀ of insulin was estimated to be between 1.7 and 2.9 nM. In comparison, clinical clamp studies report insulin-mediated suppression of hepatic glucose production with an EC₅₀ of approximately 0.15 nM for healthy individuals and 0.4 nM in patients with type 2 diabetes (30,31). This shift in potency may reflect in vitro–in vivo differences, methodological differences, variations in insulin resistance, or a combination of these factors.

Second, we observed significant insulin-independent glucose utilisation, with glucose concentrations decreasing from 11.5 mM to 7.5 mM over 48 h in the absence of insulin (**Fig. 3A a** blue). The fraction of insulin-independent hepatic glucose utilisation relative to maximal utilisation was estimated at 55%, comparable to whole-body clinical estimates (approximately 50% in non-diabetic individuals and 70% in diabetic individuals) (32,33).

Third, hepatic insulin sensitivity was found to decrease over time, as indicated by the diminishing difference between glucose uptake in insulin-stimulated conditions (**Fig. 3A c**, non-blue) and non-insulin conditions (**Fig. 3A c**, blue). The EC₅₀ of hepatic insulin-stimulated glucose utilisation increased approximately 4-fold over 15 days, resulting in the largest reductions in insulin response in groups with insulin exposures closest to the EC₅₀ (**Fig. 3A d** green).

Finally, hepatic glucose utilisation was evaluated across cultures that differed in medium volume and total cell number (**Fig. 3B**). Our results show that the cell-to-medium ratio alone did not explain the observed changes in glucose concentrations. Specifically, a 4-fold increase of this ratio in the control group resulted in an approximately 2.8-fold increase in glucose utilisation at 48 h (**Fig. 3B a** blue, red). A similar pattern emerged in cultures with identical cell-to-medium ratios: reducing medium volume twofold versus increasing cell number twofold yielded approximately 1.4-fold lower glucose concentrations (**Fig. 3B a** green, yellow). Together, these observations along with all data used to test our hypothesis, were best captured by a logarithmic scaling function in which total metabolic rates scale with total cell number with a slope of 0.45–0.63, and total intracellular volume scales with a slope of 1 (**Fig. 3B b**). These slopes align with values commonly assumed in *in vitro–in vivo* and animal–human extrapolations (34–37).

Overall, the mathematical model was able to adequately represent all data on glucose utilisation of hepatic monocultures.

### Glucose-insulin interactions in pancreatic-hepatic co-cultures

Co-culture MPS studies were designed to investigate bidirectional crosstalk between liver and pancreas, focusing on time-dependent coupling of insulin secretion and glucose utilisation, pharmacological responses to exenatide, and differences between co-cultures containing human islets versus EndoC-βH5 spheroids (**Fig. 4**).

**Figure 4:**
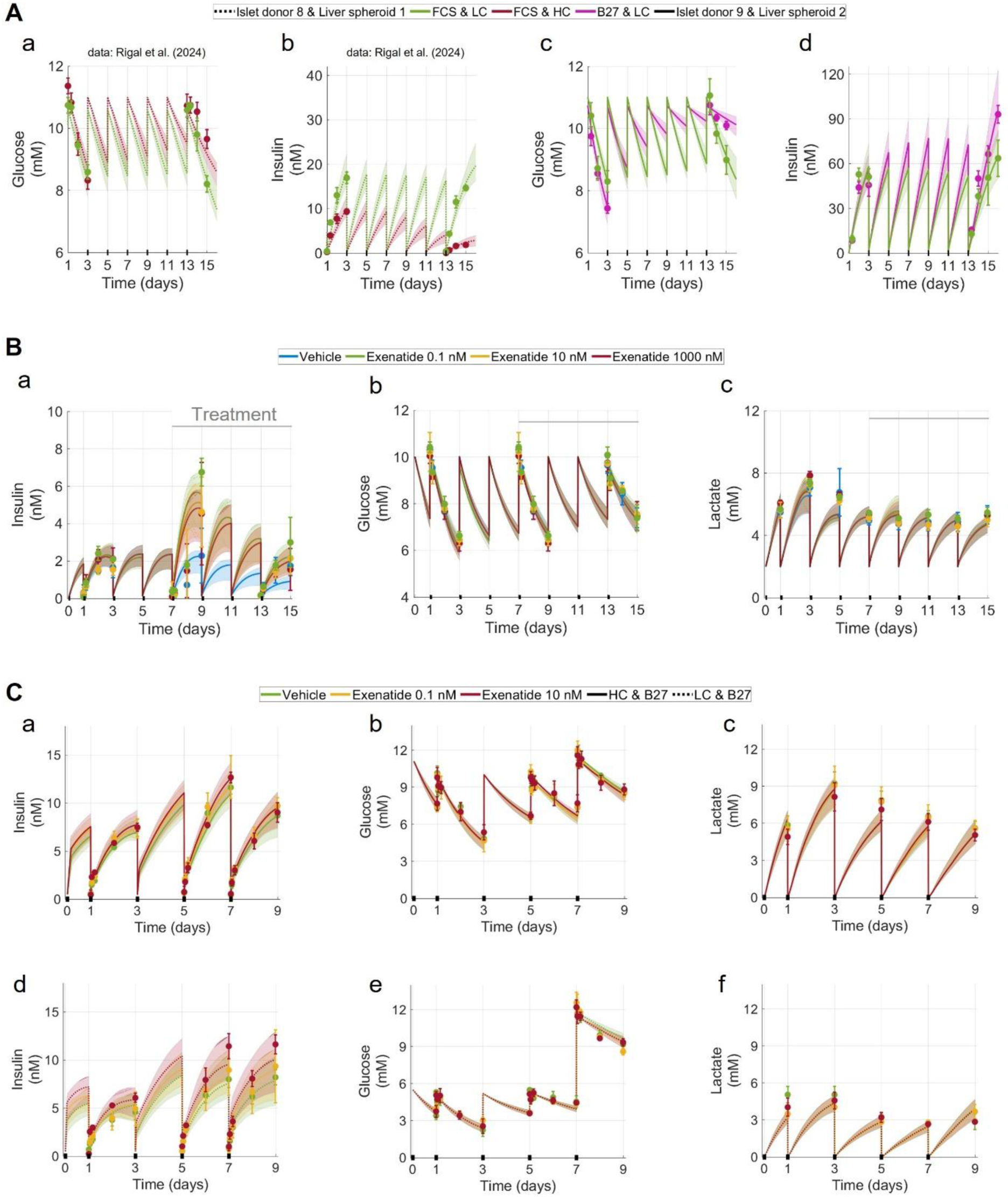
**Glucose-insulin interactions of pancreatic-hepatic co-cultures** Lines represent mathematical model best joint-fit to all cell-culture studies (Figs. 2-5, Supplementary Fig. S1). Shaded areas indicate model uncertainty. Error-bars represent data (mean ± SD). Studies were conducted with various media conditions: low hydrocortisone (LC), high hydrocortisone (HC), fetal calf serum (FCS) B27 supplements (B27). *(A)* MPS co-cultures (liver spheroids and human islet microtissues) without exenatide treatment. Dotted lines represent a previously reported study (4) (Study 1). Solid lines represent new study (Study 3). Colours represent media conditions: FCS and low cortisone (LC; green); FCS and high cortisone (HC; red); B27 and LC (magenta). a and c) Glucose in media; b and d) Insulin in media. *(B)* MPS co-culture (liver spheroids and human islet microtissues) with exenatide treatment (Study 4). Grey lines indicate duration of exenatide treatment. a) Insulin in media. b) Glucose in media. c) Lactate in media. *(C)* MPS co-culture (liver spheroids and EndoC-βH5 spheroids) with exenatide treatment (Study 11). (a and d) Insulin in media. (b and e) Glucose in media. (c and f) Lactate in media.

First, differences in insulin secretion and glucose utilisation were observed in two human islet co-cultures maintained under three media conditions (**Fig. 4A**). Across all cultures, there was a time-dependent functional decline in cell model metabolic rates, including reductions in glucose utilisation (e.g. **Fig. 4A a**, red), insulin secretion (e.g. **Fig. 4A b**), and insulin clearance (e.g. **Fig. 4A c-d**). The rate of decline and the functional baseline level both varied across media conditions (**Fig. 4A**, colours) and between cell batches and donors (**Fig. 4A**, dotted or full lines).

Second, the effect of exenatide was assessed in human islet co-cultures, where treatment initiated seven days after study onset increased pancreatic insulin secretion by approximately 3-fold (**Fig. 4B a**, green). The resulting dose-response indicated a pronounced bell-shaped relationship, with a maximal effect at 0.1 nM exenatide. The magnitude of the decline in exenatide’s effect beyond EC_100_ has been reported in previous studies (27), although not consistently in all reports, potentially because the reduction in effect may only become apparent when concentrations exceed the therapeutic range and typical clinical dosing levels (2,38,39). A peak response at 0.1 nM exenatide was repeated in static cell assays using the same human donor (Supplementary **Fig. S1E**). No significant difference was observed in end-point glucose concentrations between day 3 and day 9 (**Fig. 4B b**), suggesting that exenatide-induced increase in insulin concentrations only marginally restored glucose utilisation (**Fig. 4B b**, day 9). Restoration of glucose utilisation under pre-diabetic conditions is consistent with expected clinical responses to exenatide (40,41).

Finally, two EndoC-βH5 spheroid co-cultures under different media conditions were conducted to compare them with human islet co-cultures (**Fig. 4C**). In contrast to human islet co-cultures, EndoC-βH5 spheroids exhibited rapid insulin secretion immediately after each medium replacement, followed by a lower, sustained secretion phase (**Figs. 4C a** and **4C d**). In the mathematical model, this behaviour is explained by a biphasic insulin-secretion process. The first phase (phase I) is characterised by rapid secretion of preformed intracellular insulin stores, and an insulin production dependence on the rate of change of intracellular glucose. The second phase (phase II) is characterised by a slower insulin production that depends on intracellular glucose concentration. The output of this biphasic insulin secretion is consistent with clinical observations (42,43). An exenatide response was observed predominantly within the first two hours of the EndoC-βH5 co-cultures, and was greater under low-versus high-hydrocortisone conditions (**Figs. 4C a** and **d**). Similar insulin concentrations were observed between the two EndoC-βH5 spheroid co-cultures, suggesting that the differences in media conditions equalised the differences derived from differences in glucose concentrations (**Figs. 4C**, dotted line and full line). We found that the same mathematical representation of phase I/phase II insulin secretion that adequately described the EndoC-βH5 data could also describe insulin secretion in human islets from two preexisting studies (**Fig. 5**) (23,24).

**Figure 5:**
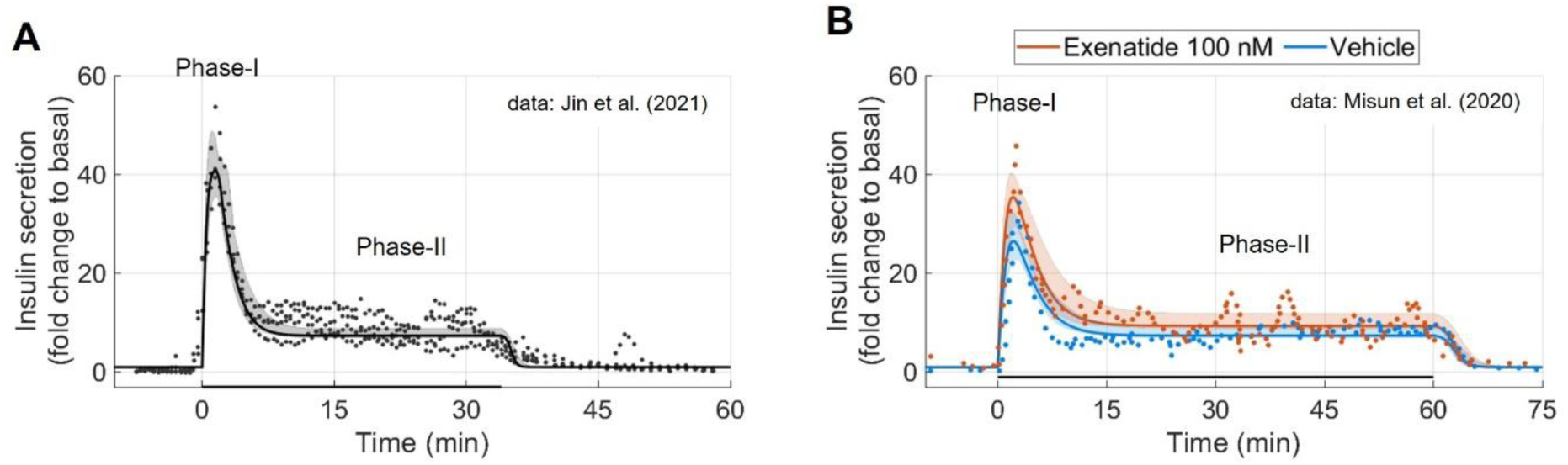
**High-time resolution insulin secretion** Lines represent mathematical model best joint-fit to all cell-culture studies (Figs. 2-5, Supplementary Fig. S1). Shaded areas indicate model uncertainty. Circle-markers are reported individual data-points. *(A)* Human islet insulin secretion (Study 18) (23). Y-axis is normalised to study mean basal insulin secretion. *(B)* Human islet insulin secretion (Study 17) (24). Y-axis is normalised to study mean basal insulin secretion.

Overall, the mathematical model was able to adequately represent all data on pancreatic-hepatic co-cultures.

### Sources of inter-study differences

We established a translation methodology across the human cell culture studies (**Figs. 2-5, Supplementary Fig. S1**), enabling all datasets to be described with a unified conclusion (**Fig. 6**). The translation methodology incorporates condition-specific parameter adjustments (condition alterations), to capture differences from a general population (e.g., all human donors) to a specific condition (e.g., human donors in a specific medium).

**Figure 6:**
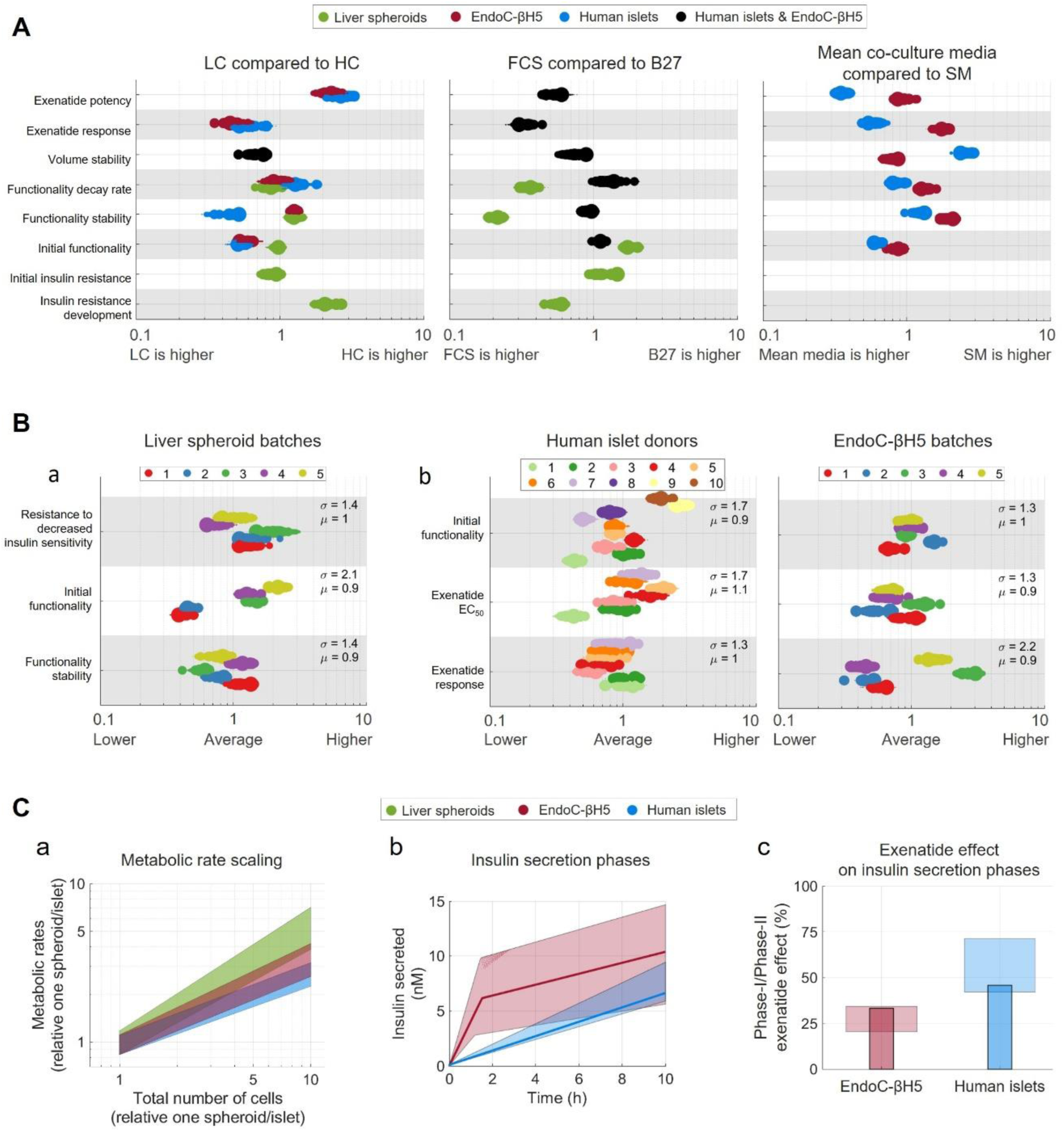
**Differences between experimental conditions** Data-driven insights generated from computer simulations with non-rejected agreement to all cell-culture studies (**Figs. 2-5**, **Supplementary Fig. S1**). Bars, dots, and lines are best agreement to data. Shaded areas indicate model uncertainty. *(A)* Media condition comparisons: low hydrocortisone (LC), high hydrocortisone (HC), fetal calf serum (FCS), B27 supplements (B27), and suppliers’ media (SM). The comparisons are made over the x-axis indicating how media conditions affect metabolic reactions or variables (y-axis). *(B)* Human islet donor and pancreatic β cell-line batch comparisons. The comparisons are made over x-axis indicating how the donor and batch differences affect metabolic reactions (y-axis). Population mean (µ) and SD (σ). a) Liver spheroid batch variability. b) Pancreatic variability: Human islet donors (left); EndoC-βH5 batches (right). EndoC-βH5 batch 1-4 were studied in spheroids and batch 5 was studied in monolayers. *(C)* Described differences between all tested cell-types. a) Metabolic rate scaling relative one spheroid/islet. b) Insulin secretion at media changes. c) Exenatide effect on the two phases of insulin secretion.

Condition alterations were used to describe media effects (**Fig. 6A**). Three comparisons were made: co-culture media with low hydrocortisone (LC) versus high hydrocortisone (HC) (**Fig. 6A** left); co-culture media with fetal calf serum (FCS) versus B27 supplement (B27) (**Fig. 6A** middle); and mean co-culture media versus suppliers’ medium (SM) (**Fig. 6A** right).

We found sufficient agreement with data when assuming identical effects of FCS and B27 on the pancreatic cell models (**Fig. 6A** black). In contrast, the same assumption was rejected for hydrocortisone levels (**Fig. 6A left** blue and red), where human islets maintained stable insulin secretion for a longer time in LC compared to HC (**Fig. 6A** blue, approximately 2.5-fold) and no such hydrocortisone-dependent effect could be concluded for the EndoC-βH5 cultures. For the liver spheroids, HC accelerated insulin resistance development relative to LC (**Fig. 6A** green approximately 2-fold). Additionally, accelerated insulin resistance development was also identified from elevated intracellular glucose, as all accepted model solutions displayed insulin resistance development above 4 mM (Supplementary **Fig. S1D**). LC compared to HC were best described with higher exenatide efficacy (approximately 2-fold), lower exenatide potency (approximately 2.5-fold), and higher cell proliferation (approximately 1.5-fold). In the mathematical model, cell proliferation was allowed to be affected by exenatide exposure, where no significant exenatide effect could be concluded (Supplementary **Fig. S1A**).

For FCS versus B27 media alternatives (**Fig. 6A**, middle), FCS delayed functional decline in liver spheroids (approximately 5-fold), which could not be concluded in the pancreatic models. FCS was also characterised with higher exenatide efficacy (approximately 2.5-fold) and may have induced insulin resistance more rapidly than B27 in liver spheroids (approximately 1.5-fold).

For the mean co-culture media versus SM (**Fig. 6A**, right), SM produced a slightly slower functional decline in EndoC-βH5 cultures (approximately 1.5-fold), while no clear difference was observed for human islets (confidence intervals are overlapping x = 1). In human islets, mean co-culture media yielded higher exenatide potency and efficacy (approximately 3-fold and 2-fold, respectively), whereas EndoC-βH5 cultures showed higher efficacy in SM (approximately 1.5-fold).

Condition alterations were also used to evaluate human donor-to-donor and cell line batch-to-batch variability (**Fig. 6B**). The condition alterations providing the best agreement with the data were used to estimate overall population mean (**Fig. 6B** µ) and population variability (**Fig. 6B** σ). All estimated population means (µ) ranged from 0.9 to 1.1, indicating sufficient containment of parameter variability. The largest population variabilities were observed in liver spheroid baseline functionality (σ = 2.1) and EndoC-βH5 exenatide efficacy (σ = 2.2). Human donor 1 exhibited lowest exenatide potency compared to all other donors, and EndoC-βH5 batch 3 displayed highest exenatide efficacy compared to all other batches (discussed in **Supplementary Note 1**).

One final remark of inter-study variation concerns differences between EndoC-βH5 cultures, human islets, and liver spheroids (**Fig. 6C**). We found that the same logarithmic scaling function used to relate liver spheroid metabolic rates to total cell number also described the metabolic rates of the pancreatic models (**Fig. 6C a**; Liver spheroids: 0.55 to 0.79; Human islets: 0.38 to 0.54; EndoC-βH5 cultures 0.45 to 0.63). Furthermore, data indicated that EndoC-βH5 cultures were more dependent on a burst of insulin secretion after media changes compared to human islets (**Fig. 6C b**), a pattern that is visually evident in the MPS studies (**Figs. 4A–B** compared to **Fig. 4C**). Finally, we found that exenatide affects both phases of insulin secretion (**Fig. 6C c**), with human islets best described by a slightly greater effect on the first phase compared with EndoC-βH5 cultures (**Fig. 6C c** blue is higher than red).

### Validation of described biology through its ability to predict new independent data

To validate the mathematical model representation of the cell cultures, we predicted the effects of exenatide with qualitative agreement to general pancreatic metabolism and quantitative agreement to new independent data (**Fig. 7**).

**Figure 7:**
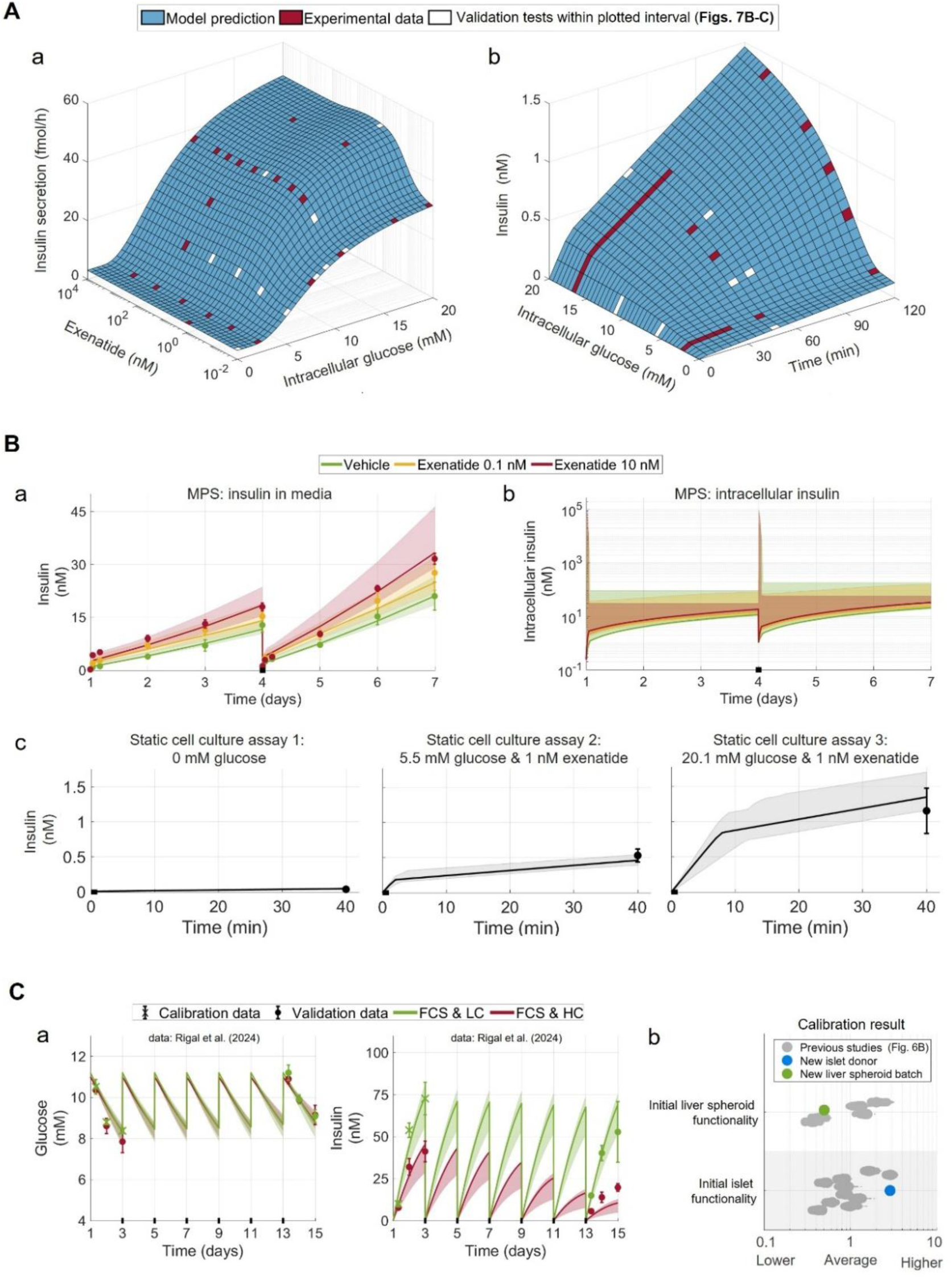
**Validation of the hypothesis through its ability to predict new independent data** Lines represent best model prediction from joint-fit description of all cell culture studies (**Figs. 2-5, Supplementary Fig. S1**). Shaded areas indicate model uncertainty. Error-bars represent data (mean ± SD). *(A)* Predicted insulin secretion dependency to time, intracellular glucose concentration, and exenatide exposures. Squares are model predictions using parameters with best agreement to all estimation data. a) insulin secretion (y-axis) dependency to exenatide (z-axis) and glucose (x-axis). b) insulin in media (y-axis) dependency to glucose (z-axis), and time (x-axis). *(B)* Predictions of EndoC-βH5 spheroid insulin secretion. Area represents model uncertainty. a) Predicted insulin in media of monoculture MPS study (Study 9). b) Predicted intracellular insulin of monoculture MPS study (Study 9). c) Predicted static cell culture assays (Study 10). *(C)* Predicted MPS co-culture (Study 2) (4). Area represents model uncertainty. a) Glucose in media (left) and insulin in media (right). Error-bars marked with a cross are calibration data. b) Calibration result. Grey dots show islets and liver spheroids present in estimation data (Fig. 6B) and coloured dots is new calibration result to the new co-culture (red and blue).

The mathematical model illustrates exenatide’s effect on insulin secretion (**Fig. 7A**). The illustration goes beyond any singular cell culture study, as it covers all studies evaluated (**Fig. 7A**, red) and provides mechanistically based estimates for conditions not measured (**Fig. 7A**, blue). The model reflects *in vitro* biology consistent with pancreatic metabolism and clinical responses: a basal insulin secretion (44), a sigmoidal glucose- and bell-shaped exenatide-dependent insulin secretion (25,26), a minimum threshold for glucose-dependent insulin secretion (29), exenatide affected glucose-dependent insulin secretion (28), and a time-dependent biphasic insulin secretion (42,43). Note that not all parts of the modelled *in vitro* insulin secretion are represented in the visualisation (**Fig. 7A**), as insulin secretion is also described to be influenced by long-term changes (e.g., secretion capacity) and experimental factors (e.g., cell type, donor or cell-line batch, media conditions and volume, and total cell number). Simulation settings were set in accordance with the standardised GSIS protocol (method section and **Supplementary note 1**).

For further validation, three independent studies were predicted: two EndoC-βH5 studies (**Fig. 7B**) and one co-culture of human islet and liver spheroids (**Fig. 7C**). These datasets were not used to train the model. The predictions passed a χ² goodness-of-fit test confirming acceptable fit (64 < 83 = 𝑇^𝑜^ = ℱ^cdf−inv^(0.95,63)).

In the first validation study, EndoC-βH5 spheroid monocultures were studied in the liver-pancreas MPS platform (**Fig. 7B a**). The mathematical model successfully predicted three key observations: a) appropriate glucose-dependent insulin secretion at 5.5 mM glucose (days 0–3) and 11 mM glucose (days 3–6); b) rapid phase-I insulin secretion following medium changes; and c) the correct dose-response relationship between the two exenatide exposure groups, where the 10 nM response exceeded the 0.1 nM response. This dose-response prediction was non-trivial, as some studies used to train the model displayed higher responses at 0.1 nM than at 10 nM (**Fig. 4B a** and Supplementary **Fig. S1E**). Furthermore, the prediction includes an accumulation of intracellular insulin stores during phase-II insulin secretion (**Fig. 7B b**), which is consistent with clinical observations (45,46).

In the second validation study, outcomes from three static cell assays of single EndoC-βH5 spheroids were successfully predicted (**Fig. 7B c**). These predictions indicate both phase-I and phase-II insulin secretion and correctly described the scaling of insulin secretion between the MPS study (**Figs. 7B a** and **b**) and the static cell assays (**Fig. 7B c**), despite a 10-fold difference in spheroid number.

In the third validation study, a new human islet donor and a new liver spheroid batch were studied in MPS (**Fig. 7C**). To calibrate for the new co-culture a minimalistic approach was chosen, where two condition alterations describing initial liver spheroid and human islet functionality were calibrated to describe the first media change of the culture with FCS and LC (**Fig. 7C a,** green; calibration data marked ‘x’). With the minimalistic calibration, the glucose and insulin concentrations of the final media change were successfully predicted (**Fig. 7C a,** validation data). Most noticeably, the description of media impact on insulin secretion was validated, where FCS and HC (**Fig. 7C a,** red) provided an earlier reduction in insulin secretion and lower insulin secretion stability compared to the FCS and LC (**Fig. 7C a,** green). The calibration result was compared to the populations used in the model training without any unplausible condition alteration outliers (**Fig. 7C b**).

Thus, arguably for the first time within a single framework, we have described *in vitro* liver-pancreas metabolism in response to a GLP-1RA and linked it to clinical observations (**Figs. 2-5**), uncovered new insights of previously unmodeled sources of inter-study variability (**Fig. 6**), and integrated all information to generate accurate drug predictions (**Fig. 7**).

## DISCUSSION

In this study, we present and interpret new data on liver–pancreas metabolism using a novel pragmatic approach suitable for heterogeneous *in vitro* datasets (**Fig. 1**). The strength of our approach lies in its ability to account for inter-study differences, enabling unified conclusions across all studies. We demonstrate that our hypothesis describes all available data: insulin secretion in pancreatic mono-culture studies (**Fig. 2**), hepatic glucose utilisation in liver spheroid monocultures (**Fig. 3**), glucose-insulin interplay in pancreatic-liver co-cultures (**Fig. 4**), and pre-existing high-time-resolution insulin secretion studies (**Fig. 5**). Importantly, we predicted previously unquantified sources of inter-study variability (**Fig. 6**), including cell-type–specific interactions with media conditions (**Fig. 6A**) and differences among cell donors and batches (**Fig. 6B**). Finally, the hypothesis was validated by successfully predicting three new independent studies (**Fig. 7**), demonstrating predictive capability in MPS (**Fig. 7B**; top), static cell cultures (**Fig. 7B**; bottom), and new liver-islet co-cultures (**Fig. 7C**). Using our approach, we identified multiple aspects of in vitro pancreas–liver biology and study-related variability. Some biological aspects aligned with clinical observations, such as the exenatide effect on a biphasic insulin secretion (**Fig. 7A**), whereas others diverged, including lower hepatic insulin sensitivity (**Fig. 3A**) and a time-dependent decline in metabolic rates in cell cultures (**Fig. 4A**). Study-specific differences were also resolved, including medium-related effects such as accelerated development of insulin resistance under high hydrocortisone and elevated glucose, and donors and batch effects, such as human donor 1 exhibiting the highest apparent exenatide potency (**Fig. 6B**). Collectively, these findings demonstrate, to our knowledge for the first time, that heterogeneous *in vitro* datasets can be integrated within a single mathematical-modelling-framework to generate unified conclusions about drug effects and *in vitro* biology.

Testing a single hypothesis against all available cell studies was made possible by our pragmatic approach in a similar vein as traditional population-based modelling. This approach introduces condition alterations, which should be evaluated in three ways for a rigorous implementation. First, the condition alterations should be assessed for necessity to describe data. We address this by penalising them so that unnecessary alterations collapse to no change (i.e., x = 1), as observed for several conditions in our analysis (**Fig. 6**). Second, the implementation requires a balance between allowed and restricted variability. This is done by introducing and evaluating a penalty-term (**Eq. 5**; Methods). Excessive penalization can override true population differences embedded in the data, leading to poor model performance. In contrast, insufficient penalization limits information integration across studies and increases the risk of overfitting to experimental noise. We demonstrate an appropriate balance, evidenced by identifiable condition alterations (**Fig. 6**), cases where alterations collapse to no change (**Fig. 6**, overlapping x = 1), good agreement with estimation data (**Figs. 2–5, Supplementary Fig. 1**), and sufficient predictions of independent data (**Fig. 7**). Third, differences in condition alterations must be evaluated for statistical significance. This is assessed by examining overlapping parameter uncertainties. To further test significance, additional penalty terms that minimize differences between condition alterations describing the same metabolic function can be introduced (**Supplementary Note 1**). Using this approach, we identified only one significant difference in how hydrocortisone affects the two pancreatic cell models (**Fig. 6A**, left; blue and red). Together, these three evaluation steps, combined with validation tests, reduce the risk of biased inference, identifiability issues, and model misrepresentation.

Our condition alteration-based approach is a reasonable choice for jointly testing hypotheses across heterogeneous *in vitro* datasets. This is because it aligns with the structure of the data (few replicates per condition and additive effects from multiple conditions) and with our objective of insight integration across studies. In principle, a traditional population-based modelling approach could encode conditions as covariates and donors/batches as random effects, e.g., non-linear mixed effect (NLME) modelling (20,21). However, to capture multiple interacting condition effects on the same metabolic rates would require complex covariate hierarchies and correlated random effects, increasing identifiability and computational burden. In addition, NLME’s likelihood-centered inference does not naturally provide the type of χ²-based global acceptance, qualitative constraints, and explicit control over how much parameters are allowed to change. In our approach, we enforce small parameter changes and permit larger deviations when the data clearly require them, which keeps the model interpretable and aligned with mechanistic expectations. For our current type of data, we therefore prefer the penalty-based strategy for its clarity, manageable estimation with fewer parameters, and direct mechanistic attribution of variability. We still recognize that an NLME model with covariates and shrinkage could be a useful complementary option once individual-level time courses and more extensive replication are available.

Testing hypotheses against variable data (multiple measured variables, timescales, and experimental conditions) enables evaluation of advanced and knowledge-rich hypotheses. This is because the complexity of mathematical-modelled-hypothesis needs to reflect data variability to avoid unidentifiability and overfitting to experimental noise. These problems are reduced by testing a hypothesis to studies jointly, as studies are generally conducted with different aims, different measured variables, different experimental conditions, and at different timescales etc. By integrating 21 cell studies in this analysis, we obtained sufficient data variability, enabling us to further develop our previous mathematical modelling (4,8–10) to a higher level of hypothesis complexity.

Our mathematical modelling analysis relies on several assumptions that could be further refined. The most important assumptions are embedded in the choice of mechanisms and equations, and we do not claim that these mechanisms are uniquely correct. For example, alternative explanations for EndoC-βH5 culture variability may exist beyond those considered here (**Supplementary Note 1**). One potential improvement is to incorporate a dimensional structure into the metabolic scaling function, which scales total metabolic rates in an experiment to total cell number. This was not implemented in our exemplification. Instead, all estimated cell numbers were pooled into one summarised value, ignoring potential impact from 2D- or 3D-structures and how the cell number is divided into islets or spheroids. An addition of condition alteration describing differences in 2D- and 3D-cultures could improve model agreement to data, as previous research has shown differences between these in both pancreatic β-cells (47) and hepatocytes (48). As a final enhancement, the penalty-term weights (**Eq. 5**; Methods) could be selected automatically rather than manually, facilitating broader application and likely improving robustness.

In conclusion, we present a pragmatic approach for drawing unified conclusions from diverse datasets and apply it to characterize the GLP-1RA exenatide effect on pancreas–liver metabolism using 16 new and five pre-existing cell culture studies. By jointly evaluating all data, we link in vitro biology to clinical observations and quantify previously uncharacterized sources of inter-study variability. Our work provides new insights into preclinical diabetes research and offers the scientific community tools to support the development of pharmacological therapies using complex experimental cell systems.

## METHODS

### Experimental design

#### Pre-culture and spheroid formation

Liver spheroids were prepared as previously described (4,8–10). Briefly, terminally differentiated HepaRG cells were pre-cultured confluently for 3 days in pre-culture medium consisting of Williams’ medium E (P04-29050S4, PAN-Biotech, w/o glucose, w/o L-glutamine, w/o phenol red) supplemented with 10% fetal calf serum (FCS; 35-079-CV, Corning), 5.5 mM glucose (25-037-CIR, Corning), 1 nM insulin (P07-4300, PAN-Biotech), 2 mM GlutaMax (35050-061, Gibco), 50 µM hydrocortisone hemisuccinate (H4881, VWR), 50 µg/mL gentamycin sulfate (30-005-CR, Corning) and 0.25 µg/mL amphotericin B (30-003-CF, Corning). Cells were thawed in pre-culture medium containing 0.5 % DMSO which was raised to 2 % DMSO by a complete medium exchange one day after thawing. Human primary hepatic stellate cells (S00354, BioIVT, Lot PFP, P3-5) were pre-cultured for 2-3 days in Stellate Cell Medium (5301, ScienCell) supplemented with Stellate Cell Growth Supplement, 2% FBS and 1% penicillin/streptomycin. The liver spheroids were formed for 3 days by combining 24,000 HepaRG cells with 1,000 stellate cells in 384-well round bottom ultra-low attachment microplates (3830, Corning) in pre-culture medium without DMSO. After collecting liver spheroids in a 24-well low attachment plate (40 per well, 3473, Corning), the spheroids were incubated overnight in pre-culture medium without DMSO on a 3D rotator (PS-M3D; Grant-bio).

The EndoC-βH5 cell line was purchased from Human Cell Design (22). The cells were cultured in cell-suppliers ULTIβ1 medium (Human Cell Design) on β-coat-coated 6-well plates (9.5 x 10^5^ cells/well) for 4 days prior spheroid formation. Cells were detached using trypsin EDTA 0.05 % / 0.53 mM (25-053-CI, Corning) and 2,000 cells were seeded per well of either Corning® 96-well Clear Round Bottom Ultra-Low Attachment Microplates (7007, Corning) or Akura™ 96 Spheroid Microplates (InSphero) depending on the experiment. After four days of formation, the EndoC-βH5 spheroids were either maintained statically (ULTIβ1-medium or co-culture medium variants) or directly transferred to the MPS.

Human islets were purchased from InSphero and maintained as previously described (4,8–10). Briefly, the islets were cultured in the supplier’s 3D InSight Human Islet Maintenance Medium (CS-07-005-02; InSphero) according to the manufacturer’s instructions.

All cell cultures were maintained at 37 °C in a humidified atmosphere containing 5% CO₂, with routine medium changes every 2–3 days. Standard aseptic techniques were used, and cultures were regularly monitored for morphology and contamination. Media were pre-warmed to 37 °C before use.

#### Study design

The objective of the human cell studies was to investigate the effect of exenatide on liver–pancreas metabolism and to assess how intra-study differences influence study outcomes. Study-specific sample size and information (**Supplementary Data S1**) and replicate-specific data (**Supplementary Data S3-S18**) are available in supplementary material. Damaged or contaminated islets and spheroids were identified visually and excluded (**Supplementary Data S1**). Two studies without an exenatide response were excluded from the analysis and are reported in the supplementary material (**Supplementary Data S17-S18**), as most probable cause for outliers was determined to be external factors (**Supplementary Note S2).**

The pancreatic and liver MPS co-cultures were studied in a commercially-available HUMIMIC Chip2 96-well (4,8,9,49,50). The Chip2 contains two spatially separated organ compartments connected with a microfluidic channel where medium is re-circulated by an integrated micropump. In all studies each organ compartment was filled with 150 μL or 300 μL medium, the microfluidic channel contains about 5 µL medium, and the pump was set to a flow rate of 4.94 μL/min.

Before transfer to the Chip2, cell cultures were washed twice with phosphate-buffered saline (PBS), followed by a 2 h equilibration in upcoming study medium to remove pre-culture medium. After equilibration, 40 liver spheroids and 10 human islets or EndoC-βH5 spheroids were transferred to their respective culture compartment into fresh co-culture medium.

Static glucose-stimulated insulin secretion (GSIS) assays were performed to assess pancreatic function. Prior to GSIS assays, EndoC-βH5 spheroids were starved in Ulti-ST starvation medium (Human Cell Design) containing 0.5 mM glucose for 24 h. Human islets were washed twice with Krebs buffer containing 2.8 mM glucose before an equilibration in 70 µL of the same buffer for 2 h. The GSIS assay was carried out as described earlier (10). In brief, the EndoC-βH5 spheroids or human islets were washed twice with Krebs buffer before incubating the cells in 50 µL Krebs buffer containing varying glucose and exenatide concentrations. The assays were preformed over 40 min for EndoC-βH5 spheroids (22) and over 2 h for human islets (10,24).

Static assays with liver spheroids were performed with 40 or 80 liver spheroids into a 24-well plate in either 605 or 305 µL co-culture medium and maintained for 15 days. The co-culture medium, used in MPS studies as well, was composed of Williams’ medium E (11 mM glucose), 10% FCS, 2 mM L-glutamine, 50 µM hydrocortisone, 50 μg/ml gentamycin sulfate, and 0.25 μg/mL amphotericin B with varying insulin concentrations (0 nM, 5 nM, 10 nM, and 100 nM).

Samples for both MPS and static cell assays were analysed according to the manufacturer’s instructions (Glucose: 1070-500, Stanbio Laboratory; Insulin: 10-1113-10, Mercodia; Exenatide: ELISA, ab272192, Abcam). Samples were collected after incubations and stored at −80 °C until analysis. Reported data in MPS studies are the mean concentration between both organ compartments (51,52).

Media used in all cell studies were either cell supplier’s medium (Human islets: Human Islet Maintenance Medium (InSphero); EndoC-BH5: ULTIβ1-medium (Human Cell Design)) or various compositions of a Williams’ medium E supplemented with 2 mM glutamine and antibiotics (4,8–10). The compositions include three choices: a) 11 mM glucose or 5.5 mM glucose; b) 50 µM hydrocortisone (HC=high hydrocortisone), 10 nM hydrocortisone (LC=low cortisone), or no additional hydrocortisone (used in combination with B27, as this contained physiological levels of the glucocorticoid corticosterone); and c) either 10% FCS, 1% B27, or 1% BSA. Working stocks of exenatide were prepared in respective media/buffers from a 5 mM stock in low protein binding tubes (22431064, Eppendorf) and stored at -80 °C. Various concentrations of exenatide for empty chip and exposure studies were prepared freshly in low protein binding tubes (30108302, Eppendorf) on the day of use.

### Mathematical modelling and statistical analysis

The general mathematical modelling follows a data-driven hypothesis testing approach which has been described in previous works (53,54). All mathematical models presented in this work are constructed using *ordinary differential equations* (ODEs) and follow the same general formulation (**Eqs. 1A-C**).

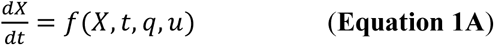

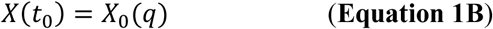

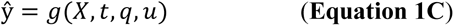

where 𝑓 and 𝑔 are non-linear smooth functions dependent on: 𝑋 which is a vector of state variables, for example concentrations of metabolites in specific compartments; 𝑞 which is a vector of model parameters, representing for example metabolic rates and scaling constants; 𝑢 which is a vector of input signals, for example media changes and drug administration; 𝑋(𝑡_0_) which is a vector of initial conditions 𝑋_0_(𝑞) that are declared from model parameters 𝑞; ŷ which is a vector of model outputs, for example metabolites in media, insulin sensitivity, etc.

The full mathematical model is reported, illustrated, and in-detail described in the supplementary material (**Supplementary Note S1**).

#### Parameter estimation

The agreement between the model simulation and experimental data was evaluated using sum weighted least squares (**Eq. 2**):

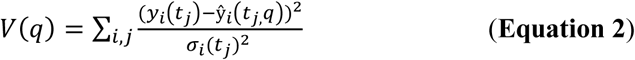

where 𝑦_𝑖_(𝑡_𝑗_) is the measured mean of a group of datapoints; where 𝑗 is a specific datapoint within the experimental setup 𝑖 at the time-point 𝑡; where 𝜎_𝑖_(𝑡_𝑗_) is the standard deviation of the group of datapoints; where ŷ_𝑖_(𝑡_𝑗,_𝑞) is the mathematical model equation value. To minimise model overfitting to small standard deviations, the mean 𝜎_𝑖_(𝑡_𝑗_) of the group was used. The weighted sum of residuals 𝑉(𝑞) was minimised in an optimisation problem to find simulations with the best agreement to data.

MATLAB R2019b (Natick, Massachusetts: The MathWorks Inc) was used for all parts of the mathematical modelling including simulations, plotting, and parameter estimations. The mathematical model was implemented in MATLAB using IQM toolbox (55). For the parameter estimation, we employed two distinct optimisation algorithms: a) particle swarm (MATLAB global optimisation toolbox); b) PESTO Monte Carlo sampling (56). The parameter estimation was done with 16 parallel workers (MATLAB Parallel Computing Toolbox) with a stopping criteria defined with max stall iterations (700), and a function tolerance (10^-5^). Parameter uncertainty was assessed from a collection of statistical non-rejected parameter sets during global and local refinement.

A statistical *χ*^2^-test was performed to quantitively evaluate a model’s agreement to data. The statistical test was done with the null hypothesis that the experimental data had been generated by the model, assuming that the experimental noise was additive, independent, and normally distributed (54). In practice, the agreement to data was rejected if the weighted residuals 𝑉(𝑞) had a higher value than the 𝜒^2^-threshold. This is given by the inverse *χ*^2^ cumulative density function (**Eq. 3**):

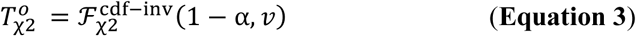

where ℱ^cdf−inv^ is the inverse density function; α is the significance level (α = 0.05 was used); where *v* is the degrees of freedom, which was equal to the number of data points in the dataset. In addition to statistical *χ*^2^-tests, model rejection was made through qualitative assessments. These qualitative assessments were implemented into the objective function where the output describes the agreement to both qualitative dynamics and data (**Eq. 4**):

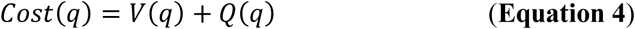

where 𝐶𝑜𝑠𝑡 is the objective function that is minimised in the parameter estimation with parameters 𝑞; where 𝑉(𝑞) is model agreement to data; where 𝑄(𝑞) is a function returning a scalar value based on the model agreement to qualitative system dynamics. Residuals of subject-specific data were taken into consideration in the qualitative assessment term due to a lack of standard deviation 𝜎_𝑖_(𝑡_𝑗_).

### Introducing condition alterations

Model parameters are structured in two layers: a) a base parameter, which describes a value applicable to all conditions (e.g. the EC_50_ value of exenatide for all human islets); and b) a condition alteration, which describes a change from the base parameter to a specific condition (e.g. the EC_50_ value of exenatide for a specific human islet donor). The difference between the base parameter and the condition-specific parameter can be called a parameter adjustment step, which is calculated in the objective function (**Eq. 5A**):

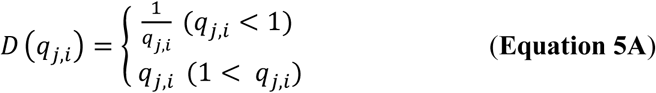

where 𝐷 (𝑞_𝑗,𝑖_) is the parameter adjustment step; where 𝑞_𝑗,𝑖_ is the condition alteration; where 𝑗 indicates which base parameter is altered; where 𝑖 specifies the condition that changes the base parameter. The adjustment step is minimised with a penalty-term in the objective function (**Eq. 5B**):

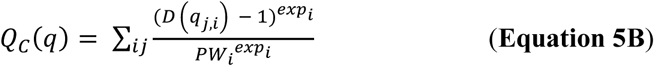

where 𝑄_𝐶_(𝑞) is a penalty term that regularizes condition-specific alterations towards 1; where 1 denotes no change to the base parameter because condition alterations act multiplicatively on the base parameter; where 𝑃𝑊_𝑖_ is the penalty weight for condition i; and where 𝑒𝑥𝑝_𝑖_ is an exponent used to scale the impact of the constraint on the objective function. The result of the minimisation is that differences between conditions are reduced, enabling the integration of insights across conditions. The parameter 𝑃𝑊_𝑖_ determines the strength of the penalty term: a high value implies that parameter estimation is primarily driven by agreement with the data, whereas a low value allows agreement with the data to be traded for increased insight translation between conditions. The parameter 𝑒𝑥𝑝_𝑖_ determines the degree of exponential penalisation in the parameter adjustment step.

Our exemplified implementation is described in detail in the supplementary material (**Supplementary Note 1**). In practice, 𝑒𝑥𝑝_𝑖_ was set to two when the penalty term was applied to condition alterations affecting the same model parameter. An exponential punishment is important when several condition alterations influence the same model parameter, as it reduces a source of parameter unidentifiability and facilitates collapsing condition alterations into one if they are not required to describe the data. A linear punishment term (𝑒𝑥𝑝_𝑖_ = 1) was used when parameter differences were described by only one parameter, in order to penalise both small and large parameter steps effectively. The manual assignment of 𝑃𝑊_𝑖_ was set to 1.5 when minimising differences between EndoC-βH5 cultures and human islets, and to 1 when describing differences between media conditions and cell donors/batches. These values were chosen to balance allowed and constrained parameter variability, so that the qualitative system dynamics 𝑄(𝑞) was approximately 10% of the total 𝐶𝑜𝑠𝑡 (**Eq. 4**).

## DATA AVAILABILITY

The analysis includes previously reported data by us (**Supplementary Data S3-S4**) (4), new studies (**Supplementary Data S5-S18**; Clinical trial number: not applicable), and data collected from literature (**Supplementary Data S19-21**) (22–24). The data previously reported by us was downloaded from the publications supplementary material (4). The data collected from literature was digitalised using a publicly available software (57). All data is available in the supplementary material and from our GitHub repository (https://github.com/OscarSilfvergren/Addressing-Study-Variations-in-Heterogeneous-Cell-Systems.git).

## CODE AVAILABILITY

All code used in the analysis is available to be downloaded from our GitHub repository (https://github.com/OscarSilfvergren/Addressing-Study-Variations-in-Heterogeneous-Cell-Systems.git).

## ACKNOWLEDGEMENTS

**Johanna Dietzfelbinger** (TissUse) for figure visualisation.

**Christopher Rhodes** (AstraZeneca) for discussions regarding insulin secretion biology. **Christine Schwenk** (InSphero) for insights in exenatide effect and pancreatic-hepatic crosstalk. **Bruno Blanchi** (Human Cell Designs) for discussion regarding EndoC-βH5 cell biology.

The **MPS network** (AstraZeneca) for feedback and discussions over the years the analysis was done.

## FUNDING

Swedish Research Council 2023–03186 (GC) Swedish Research Council 2023–05460 (GC)

X-HiDE, Knowledge Foundation 20200017 (GC) CENIIT 15.09 (GC)

The Horizon Europe project STRATIF-AI 101080875 (GC)

The Swedish Fund for Research without Animal Experiments F2019-0010 (GC) ELLIIT 2020-A12 (GC)

The County Council of Östergötland RÖ-1001928 (GC) VINNOVA VisualSweden 2020-04711 (GC)

## AUTHOR INFORMATION

### Authors and Affiliations

**Linköping University, Linköping, Sweden**

Oscar Silfvergren, Christian Simonsson, Peter Gennemark, and Gunnar Cedersund

**SUND sound medical decisions, Linköping, Sweden**

Oscar Silfvergren and Gunnar Cedersund

**TissUse GmbH, Berlin, Germany**

Sophie Rigal, Katharina Schimek, and Uwe Marx

German Federal Institute for Risk Assessment (BfR), Berlin, Germany

Sophie Rigal

Technische Universität Berlin, Berlin, Germany

Katharina Schimek

AstraZeneca, Gothenburg, Sweden

Kajsa P Kanebratt, Liisa Vilén, and Peter Gennemark

InSphero AG, Schlieren, Switzerland

Felix Forschler and Burcak Yesildag

Örebro University, Örebro, Sweden

Gunnar Cedersund

## Contributions

**Conceptualization**: OS, CS, LV, PG, GC

**Methodology**: OS, SR, KS, CS, PG, GC

**Investigation**: OS, SR, KS, KPK, FF, BY, LV, PG, GC

**Visualization**: OS, SR, KS, LV, PG, GC

## Funding acquisition**: GC**

**Project administration**: LV, PG, GC

**Supervision**: BY, UM, LV, PG, GC

**Writing – original draft**: OS, SR, KS, CS, LV, PG, GC

**Writing – review & editing**: OS, SR, KS, CS, KPK, FF, BY, UM, LV, PG, GC

Corresponding author

Correspondence to Gunnar Cedersund.

## SUPPLEMENTARY INFORMATION

**Supplementary Fig. S1:** Additional human cell data used to evaluate mathematical model

**Supplementary Fig. S2:** Two co-culture MPS studies which provided no exenatide response

**Supplementary Table S1:** Study-specific design and description

**Supplementary Table S2:** List of pre-existing studies

Supplementary Tables S3 to S21: All data used herein

**Supplementary Note S1**: Detailed description of the mathematical modelling

**Supplementary Note S2**. Supplementary discussion regarding a lack of exenatide response in two studies

**Supplementary Figure S1:**
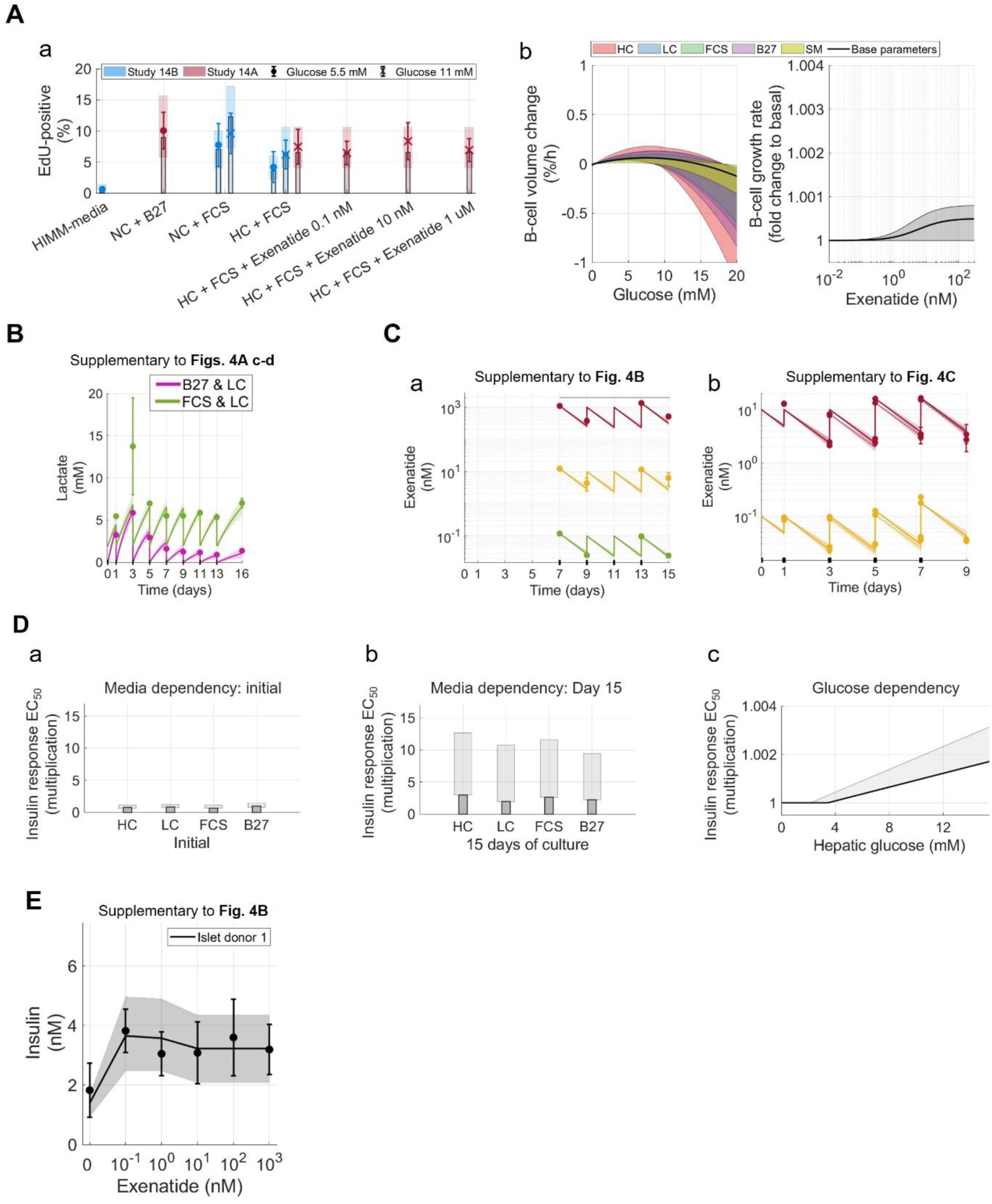
Supplementary human cell data and simulations Lines and bars represent the model’s best description of all cell culture studies together (*Figs. 2-5*, Supplementary Fig. S1). Coloured areas indicate model uncertainty. Error-bars represent population data (mean ± SD). (A) Proliferation of human islets in static cell assays (a) EdU-positive measurements (Study 14). New data is represented with the colour red (Study 14A) and previously reported data is blue (1) (Study 14B). b) Predictions of media condition and *glucose impact on cell death and growth rate. c) Rejection of significant exenatide effect on b-cell proliferation*. (B) Supplementary lactate readouts of Study 3. (C) Supplementary exenatide pharmacokinetics of two MPS co-culture studies. a) Study 4. b) Study 11. (D) Predictions of insulin resistance development. These plots are made from data to evaluate insulin resistance development. Simulations were made in accordance with GSIS experimental protocol with further explanation in the supplementary material (Supplementary Note S1) a) Initial EC50. b) EC50 after 15 days. c) Glucose-dependent insulin resistance development. (E) Human islet exenatide dose-response with 16.8 mM glucose (Study 5).

**Supplementary Figure S2.**
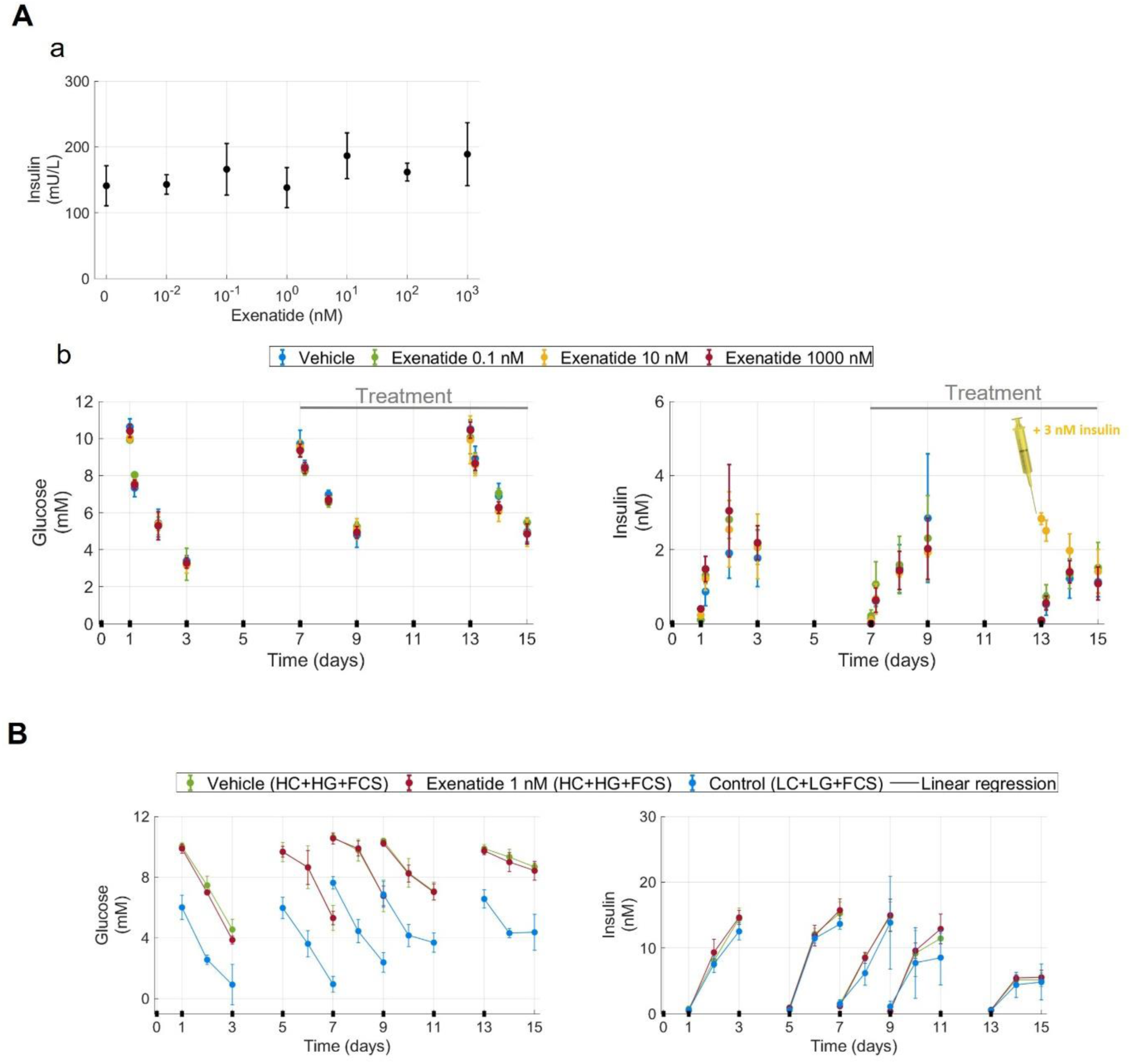
: Supplementary data of Study 15 and 16 Supplementary plot of Study 15 and 16, showing no significant exenatide response. Error bars represent population data (mean ± SD). This supplementary plot is discussed in the supplementary material (Supplementary Note S2). Data is included in the supplementary material (Supplementary Tables S17-S18). (A) Study 15. a) Static human islet assays. b) MPS co-culture study with liver spheroids. Study was done with 4 different exenatide exposures: 0 nM (blue), 0.1 nM (green), 10 nM (yellow), and 1000 nM (red). Exenatide treatment was started at day 7 (grey line, marked with treatment). At the media change of day 13, the 10 nM exenatide group got an extra insulin injection of 3 nM (yellow; marked with yellow syringe). (B) Study 16. Study was done with 3 different conditions: Vehicle (green), no exenatide, high glucose (HG), high hydrocortisone (HC), and fetal calf serum (FCS); Exenatide 0.1 nM (red), HG, HC, and FCS; and Control (blue), no *exenatide, low glucose (LG), low hydrocortisone (LC), and FCS. To increase readability of the figure, linear regression was used in between data-points of the same media changes (lines)*.

