## Supplementary Note S1 for "An Integrated Analysis of GLP-1R Agonist Mechanisms: Addressing Study Variations in Heterogeneous Cell Systems"

This supplementary note provides additional information regarding the mathematical modelling. First, a description of the usage of the model will be explained, followed by a detailed explanation of the mathematical model structure.

### Model explanation and description

The mathematical model was used throughout this study to extract additional information from the cell-culture studies than the study readouts.

The mathematical model includes parameters to describe a minimalistic glucose-central metabolism, and differences between experimental conditions. As described in the main text, the general description of the metabolism is done with base parameters (e.g. maximum exenatide response of human islets), and condition alterations which alter the parameters from describing a general population to a specific condition (e.g. maximum exenatide response in a human islet donor). The hepatic glucose metabolism was studied using liver spheroids and were described with 17 different base parameters. The pancreatic glucose metabolism (studied using EndoC-βH5 cultures and human islets) was described with 26 different base parameters. Of these 26 pancreatic parameters, 21 have two copies – one describing an EndoC-βH5 specific pancreatic metabolism and one describing a human islet specific pancreatic metabolism. These two copies of the same parameter describe the same pancreatic function. The difference between the two parameter copies was minimised in the parameter estimation to allow integration of insights across them. This was done using the equation declared in the main text regarding minimizing a parameter adjustment step (**Eq. 5B**), with the settings $PW_i$ = 1.5 and $exp_i$ = 1. The other 5 of 26 base pancreatic parameters which described both the EndoC-βH5 and human islet populations are the following: human islet cell growth rate in relation to exenatide exposure (2 parameters) and cell growth rate relation to glucose exposure (3 parameters). This means that in the mathematical model the pancreatic cell growth rate and death rate relation to glucose and exenatide are assumed to be identical (**Supplementary Fig. S1A**). This was done to minimise model complexity, as no data were found to reject the assumption. In the mathematical model, no parameter pairs exist describing the liver metabolism, as only one liver cell model was used.

As described in the main text, the impact of media conditions on the cells is described using condition alterations. This was performed by assuming that differences between studies with different media conditions can be explained by variability in pre-defined metabolic reactions (**Fig. 6A**). For the liver spheroids, these metabolic reactions were assumed to be the following: a) time-dependent loss of cell functionality, in terms of maximum functionality, functionality stability, and rate of functionality loss (3 parameters); b) initial insulin resistance (1 parameter); and c) time-dependent insulin resistance development (1 parameter). The term ‘cell functionality’ is used throughout this work and refers to total islet or spheroid metabolic rates. This ‘cell functionality’ affects all rates, e.g. glucose utilisation and insulin production. For the pancreatic cells (EndoC-βH5 cultures and human islets), the corresponding metabolic reactions were assumed to be: a) time-dependent loss of cell functionality, in terms of maximum functionality, functionality stability, and rate of functionality loss (3 parameters); b) exenatide’s potency ($EC_{100}$) and exenatide maximum response amplitude (2 parameters); and c) volume volatility (1 parameter). Exenatide's potency changes ($EC_{100}$) were assumed to be both on the positive ($EC_{50}$) and negative term ($IC_{50}$) of the bell-shaped dose-response curve. The parameter adjustment steps were minimised with the settings $PW_i$ = 1 and $exp_i$ = 2 (**Eq. 5B**). The media conditions impact on various metabolic rates and reactions was minimised with the settings $PW_i$ = 1 and $exp_i$ = 1 (**Eq. 5B**), to allow insight integration between studies.

The three cell-types used in this analysis (liver spheroids, human islets, and EndoC-βH5) were also assigned condition alterations to account for differences between donors and batches. This was done under the assumption that differences between the batches and donors could be explained by variability in pre-determined metabolic reactions. For the hepatic liver spheroids (5 batches), the differences were assumed to be the following: a) resistance to decreased insulin sensitivity (1 parameter); and b) time-dependent cell functionality, in terms of initial functionality and stability (2 parameters) (**Fig. 6B a**). For the pancreatic cultures (5 EndoC-βH5 batches and 10 human islet donors) the differences between batches and donors were assumed to be: a) exenatide's potency ($EC_{100}$) and maximum response amplitude (2 parameters); and b) initial cell functionality (1 parameter) (**Fig. 6B b**). Exenatide's potency changes ($EC_{100}$) were assumed to be both on the positive ($EC_{50}$) and negative term ($IC_{50}$) of the bell-shaped dose-response curve. The batch and donor condition alterations were minimised with the settings $PW_i$ = 1 and $exp_c$ = 2 (**Eq. 6B**), to allow insight integration between studies.

One exenatide dose-response test of EndoC-βH5 spheroids (batch 3) provided unusually low insulin secretion at low exenatide exposures (**Fig. 2A**, green). The spheroid batch had not previously displayed unusually low insulin secretion, where it was used in a glucose dose-response test (**Fig. 2E**, green). In the glucose dose-response test the culture with 11.1 mM glucose resulted in approximately 0.4 nM insulin (**Fig. 2E,** green). When the same condition was tested in the exenatide dose-response test (11.1 mM glucose and no exenatide) the resulting insulin concentration in the media was significantly lower (**Fig. 2A,** green; approximately 0.1 nM insulin). As exenatide dosages were increased in the exenatide dose-response test, the spheroids displayed an unusually high maximum exenatide response amplitude. At the final exenatide dose-response test (11.1 mM glucose and 100 nM exenatide) the GSIS result was approximately 0.6 nM insulin, which is closer to the expected value considering its glucose dose-response test (**Fig. 2E,** green; 0.6 nM is higher than 0.4 nM) and other exenatide dose-response tests (**Fig. 2A**, blue). However, we could not reject the plausibility that the spheroid batch 3 did have an unusually high exenatide response amplitude. Therefore, an assumption was made that an unspecified external factor reduced all insulin secretion of the batch 3 cultures in the exenatide dose-response study. This external factor was implemented in the model by lowering the spheroid functionality in between the glucose dose-response test and the exenatide dose-response test. This was performed in the mathematical model using the parameter 'ExperimentalInducedStress'. As described in the main text, reducing spheroid functionality means that all metabolic rates in the spheroid are reduced (e.g. glucose uptake and insulin secretion). The result of this assumption can be seen in many parts of our analysis (e.g. **Fig. 6B b,** green), where the batch ID 3 displays the highest exenatide response amplitude of all the batches.

In our analysis three plots were constructed to compare differences between media conditions and pancreatic cultures. These simulations require declared simulation settings (i.e. models inputs) which describe media volume, total cell numbers, media composition etc. Below all these model inputs are explained:

1. Four model simulations were made to plot changes in insulin resistance in various media conditions (**Supplementary Fig. S1D**). The simulations were made with the following conditions: the standardised 40 liver spheroids, no pancreatic cells, the standardised 305µL total media volume, and the standardised media change frequency of once every two days. In all conditions the media changes were simulated with 11 mM glucose, which is the standardised high glucose concentration used in all our studies (referred to HG or "high glucose" in the study protocol descriptions).

2. Two model simulations were made to illustrate EndoC-βH5 and human islets different dependency on phase-I insulin secretion (**Fig 6C b**). The simulations were made with: no liver spheroids, the standardised 10 pancreatic islets or spheroids per study, average human donor and spheroid batch, 605µL total media volume, 10 nM exenatide in media, media condition as FCS with high hydrocortisone, and one media change in the beginning of the plot. The media change was simulated to be changed from the standardised normoglycemic condition (5 mM) to hyperglycaemic condition (11 mM).

3. The mathematical model visualisation of insulin secretion dependency on time, glucose, and exenatide was made with many simulations (**Fig. 7A**), where each square on Fig. 7A a is one simulation, and every change in glucose on Fig. 7A b requires new simulations. Apart from the declared changes in glucose and exenatide, all these simulations were made with the identical GSIS settings used in the GSIS validation (**Fig. 7B c**), which was one EndoC-βH5 spheroid of batch 1 in 50 µL total media volume. The first visualisation plot shows exenatide and glucose dependent insulin secretion (**Fig. 7A a**). To make this plot the insulin secretion was plotted after 40 minutes of simulations, in agreement to the GSIS settings (40 minutes). The second visualisation plot shows time and glucose dependent insulin secretion (**Fig. 7A b**). To make this plot the insulin secretion was plotted without exenatide, to show insulin secretion independent to exenatide exposure.

All other results reported in our work were performed according to corresponding study protocols (**Supplementary Data S1**) or are direct results of parameter values and equations. These parameter values and equations are explained in detail below.

### Model structure

Each part of the model is declared and explained in four sections: a) model states, b) model parameters, c) model ordinary differential equations (ODEs), and d) model reactions.

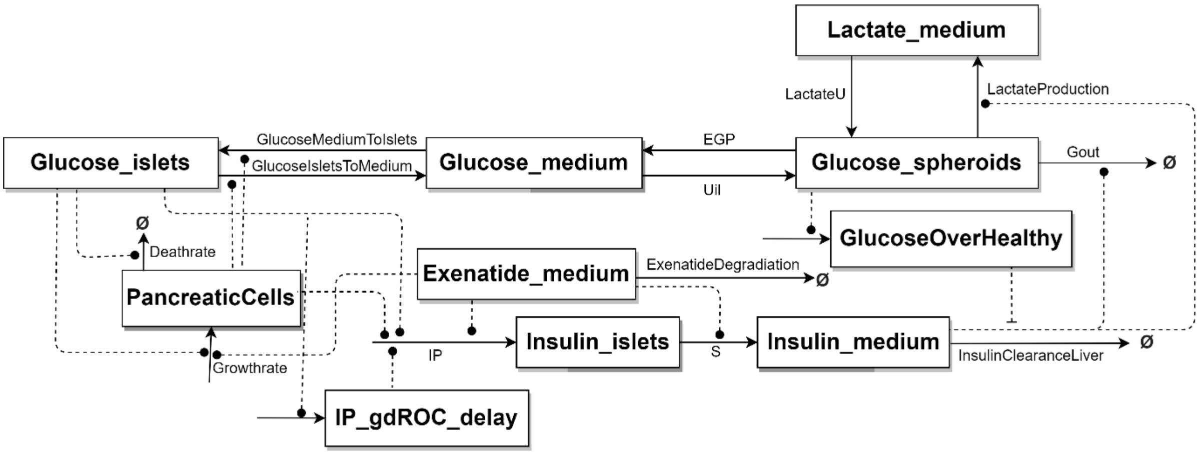


***Illustration of mathematical model***

*Rectangles represent states, text with no squares represent reactions, and arrows represent reaction rates/flows. Flows into Ø are leading out of the module. Dotted lines represent regulations, where a circle head represents a positive regulation, and a perpendicular head represents a negative regulation.*

### a) Model states

The model states are time-dependent variables which are defined by ODEs.

***List of model states***

*Θ denotes model state initial values. All model states which represent intracellular compartments are assumed from a steady state simulation (marked *), where $\theta_{EndoCLiver}$ is the initial values for liver spheroid and EndoC-βH5 co-cultures, where $\Theta_{IsletsLiver}$ is the initial values for liver spheroid and human islet co-cultures, $\theta_{Liver}$ is the initial values for liver spheroid mono-cultures, $\theta_{EndoC}$ is the initial values for EndoC-βH5 mono-cultures, and $\Theta_{Islets}$ is the initial values for human islet mono-cultures. All media compartments and number of cells in study were set in accordance with study protocol. Disease and delay state were reset between steady state simulation and study simulation (set to 0).*

| Name | Unit | Description | $\theta_{EndoCLiver}$ | $\Theta_{IsletsLiver}$ | $\theta_{Liver}$ | $\theta_{EndoC}$ | $\Theta_{Islets}$ |
|---|---|---|---|---|---|---|---|
| Glucose_islets* | mM | Intracellular pancreatic glucose | 2.56 | 2.57 | 0 | 5.50 | 5.50 |
| Insulin_islets* | pM | Intracellular pancreatic insulin | 405349.08 | 72.69 | 0 | 405349.08 | 72.69 |
| Glucose_Medium | mM | Glucose in medium | Set to media condition | | | | |
| Insulin_Medium | pM | Insulin in medium | Set to media condition | | | | |
| Exenatide_Medium | nM | Exenatide in medium | Set to media condition | | | | |
| Lactate_Medium | mM | Lactate in medium | Set to media condition | | | | |
| Glucose_spheroids* | mM | Intracellular hepatic glucose | 1.99 | 1.98 | 2.57 | 0 | 0 |
| GlucoseOverHealthy | mM | Disease state that is part of | 0 | | | | |

| | | | |
|---|---|---|---|
| | | insulin resistance development. This state sums glucose over a threshold over time. | |
| IP_gdROC_delay | nM | State that is a part of phase-I insulin production. The state introduces a delay between glucose rate of change, and its impact on insulin production | 0 |
| PancreticCells | cells | Number of total pancreatic cells in experiment | Set to experimental set-up |

The intracellular states (marked with '*' in table above) were estimated from a 48-hour steady state simulation. The media assumed in the steady state simulation was set to have 5.5 mM glucose and 1000 pM insulin, in accordance with pre-culture described in the main text.

#### b) Model parameters

The model parameters describe various metabolic rates, flows, and conversion rates. The parameters can be classified as base parameters (which describes metabolic reactions on a population-level, e.g. general human islet response to exenatide), condition alterations (which alters base parameters to condition-specific simulations, e.g. a specific human islet donor response to exenatide in a specific medium), and uncertainty parameters (which is not a part of the physiologically based model but are used to represent various uncertainties when making simulations).

These parameters are listed and described below.

***List of general parameters describing pancreatic cells.***
$\theta_{EndoC}$ *denotes the part of the parameter vector which describes the EndoC-βH5 cultures, and* $\theta_{HumanIslets}$ *denotes the part of the parameter vector which describes the human islets. For merged cells spanning both the* $\theta_{EndoC}$ *and the* $\theta_{HumanIslets}$ *columns, the same parameter is used to describe both populations.*

| Parameter | Unit | Short description | $\theta_{EndoC}$ | $\theta_{HumanIslets}$ |
|---|---|---|---|---|
| AllometricScalingK | dimensionless | Metabolic rate scaling from total cell number in experiment | 0,538 | 0,419 |
| Fislets50 | h | Time when pancreatic islets/spheroids have lost 50% of functionality | 289,604 | 371,139 |

| | | | | |
|---|---|---|---|---|
| Filets0 | fold-increase | Initial pancreatic spheroid/islet functionality | 0,969 | 0,857 |
| Filets_hill | dimensionless | Hill coefficient of pancreatic spheroid/islet loss of functionality | 3,781 | 4,056 |
| d0 | 1/h | Basal death rate | 0,001 | |
| r1 | 1/h | Glucose dependent growth rate | 0,004 | |
| r2 | 1/h | Glucose dependent death rate | 0,016 | |
| kv | dimensionless | Parameter scaling both growth and death rate | 120,415 | 4,771 |
| GlucoseIsletsMediumDiffusion | L/h | Glucose rate between pancreatic islets/spheroid and medium before functionality and allometric scaling | $1{,}11*10^{-7}$ | $1{,}11*10^{-7}$ |
| ExenatideDegradationK | 1/h | Loss of exenatide | 0,030 | |
| ExenatideInsulinMax | fold-increase | Positive term of exenatide effect on insulin secretion and production | 2,374 | 1,997 |
| Exanitide_IP | % | Exenatide effect on Phase-II/Phase-I insulin production | 0,331 | 0,457 |
| ExenatideInsulin50 | nM | Exenatide $EC_{50}$ on insulin secretion and production | 0,166 | 0,298 |
| ExenatideInsulin_hill | dimensionless | Hill coefficient of exenatide effect on insulin secretion and production | 1,332 | 4,96 |
| ExenatideInsulinDecayMax | fold-increase | Negative term of exenatide effect on insulin secretion and production | 0,377 | 0,378 |
| ExenatideInsulinDecay50 | nM | Exenatide $IC_{50}$ on insulin secretion and production | 16,119 | 16,135 |
| Exanitide_StabilisationMax | fold-increase | Exenatide effect on pancreatic growth rate | 0,001 | |
| Exanitide_StabilisationIC50 | nM | Exenatide $EC_{50}$ on pancreatic growth rate | 5,591 | |
| IP_basalK | pM/h | Basal insulin production | 0,001 | 0,002 |
| IP_gdROCk | dimensionless | Glucose rate of change (mM/h) dependent insulin production (pM/h) | 0,013 | 0,001 |
| IP_gdROC_delayK | 1/h | Delay between glucose rate of change and glucose rate of change dependent insulin production | 872,239 | 47,2 |
| IP_gdBaselineMax | dimensionless | Maximum glucose concentration (mM) dependent insulin production (pM/h) | 0,013 | 0,052 |

| | | | | |
|---|---|---|---|---|
| IP_BaselineK | mM | Glucose threshold to trigger glucose concentration dependent insulin production | 1,777 | 2,182 |
| IP_gdBaseline50 | mM | $EC_{50}$ of glucose concentration dependent insulin production | 5,372 | 5,464 |
| IP_gdBaselineHill | dimensionless | Hill coefficient of glucose concentration dependent insulin production | 1,998 | 3,881 |
| S_max | L/h | Maximum insulin secretion | 0,044 | 0,36 |
| S50 | pM | Intracellular insulin concentration where insulin secretion is 50% of maximum saturation | 373510,673 | 14815,753 |

***List of general parameters describing liver spheroids.***
$\theta_{LiverSpheroids}$ *denotes the part of the parameter vector which described the liver spheroids.*

| **Parameter** | **Unit** | **Description** | $\boldsymbol{\theta_{LiverSpheroids}}$ |
|---|---|---|---|
| ISk | dimensionless | Parameter scaling insulin sensitivity to time- and glucose-dependent functions. | 0,014 |
| IS0 | fold-increase | Initial insulin sensitivity | 0,835 |
| InsulinResistanceUpperlimit | mM | Threshold of glucose concentration triggering insulin resistance development | 3,44 |
| InsulinResistanceTimeDependency | 1/h | Time-dependent insulin resistance development | 0,258 |
| Fspheroids50 | h | Time when spheroids have lost 50 % of functionality | 218,533 |
| Fspheroids0 | fold-increase | Initial spheroid functionality | 0,982 |
| Fspheroids_hill | dimensionless | Hill coefficient of loss of spheroid functionality | 3,121 |
| InsulinResponseMax | fold-increase | Maximum insulin response | 15,903 |
| InsulinResponseLactateK | % | Ratio of glucose utilisation that becomes lactate | 0,763 |
| InsulinResponse50 | pM | Hepatic insulin response $EC_{50}$ before multiplication with diseased function | 3183,794 |
| InsulinResponse_hill | dimensionless | Hill coefficient of hepatic insulin response | 3,98 |
| Uil_iiK | L/h | Glucose rate between media and liver spheroid before functionality and allometric scaling | $2,56*10^{-5}$ |
| InsulinClearanceLiverK | L/h | Insulin clearance rate before functionality and allometric scaling | $3,00*10^{-5}$ |
| LactateProductionK | L/h | Basal lactate production rate before functionality and allometric scaling | $8,77*10^{-5}$ |
| GoutK | L/h | Basal glucose metabolization into non-lactate before functionality and allometric scaling | $1,06*10^{-6}$ |
| LactateUk | L/h | Lactate metabolised into glucose rate | 0,00016 |
| AllometricLiverScalingK | dimensionless | Metabolic rate scaling from total cell number in experiment | 0,727 |

***List of media condition alterations which affect the liver spheroids***

*The step indicates how a condition affects a base parameter. The base parameter is influenced through multiplication, where 1 means no change, 2 means doubling the parameter value, etc. The name of the condition alteration is identical to the base parameter which it affects, i.e. IS0 in the table affects the base parameter IS0.*

| Condition alteration | | **InsulinResistanceTimeDependency** | **IS0** | **Fspheroids0** | **Fspheroids50** | **Fspheroids_hill** |
|---|---|---|---|---|---|---|
| Description | | Time-dependent insulin resistance development | Initial insulin sensitivity | Initial cell functionality | Cell function stability | Cell functionality decay rate |
| Unit | | step from base parameter (fold-increase) | | | | |
| Media condition | **HC** | 1,518 | 0,948 | 0,619 | 1,372 | 0,797 |
| | **NC** | 0,738 | 1,002 | 0,633 | 1,097 | 0,918 |
| | **FCS** | 1,374 | 0,784 | 0,493 | 1,915 | 1,683 |
| | **B27** | 0,825 | 1,147 | 0,852 | 0,413 | 0,616 |

***List of media condition alterations on pancreatic cells.***

*The step indicates how a condition affects a base parameter. The base parameter is influenced through multiplication, where 1 means no change, 2 means doubling the parameter value, etc. For all condition alterations the name of the condition alteration is identical to the base parameter which it affects, i.e. kv in the table affects the base parameter kv, expect for ExenatidePotency (which affects both the negative and positive term in the potency equations: ExenatideInsulinDecay50 and ExenatideInsulin50) and ExenatideResponse (which affects both the negative and positive term in the response magnitude equations: ExenatideInsulinDecayMax and ExenatideInsulinMax). Note that EndoC-βH5 and human islets have unique relations to HC and LC condition, as described in the manuscript. For merged cells, the same parameter is used to describe the media conditions.*

| Condition alteration | | **kv** | **Exenatide Potency** | **Filets0** | **Filets_hill** | **Exenatide Response** | **Fislets50** |
|---|---|---|---|---|---|---|---|
| Description | | Impact on proliferation volatility | Exenatide $EC_{100}$ | Initial cell functionality | Cell functionality decay rate | Exenatide max and min response amplitude | Cell function stability |
| Unit | | step from base parameter (fold-increase) | | | | | |
| Media condition | $HC_{EndoC}$ | 1,601 | 0,518 | 1,08 | 1,484 | 0,73 | 0,453 |
| | $HC_{HumanIslets}$ | | 1,185 | 2,051 | 1,639 | 1,618 | 0,36 |
| | $LC_{EndoC}$ | 1,233 | 0,446 | 0,933 | 1,666 | 0,846 | 0,494 |
| | $LC_{HumanIslets}$ | | 1,184 | 1,823 | 1,3 | 1,616 | 0,951 |
| | FCS | 1,213 | 0,422 | 1,929 | 0,737 | 1,837 | 1,446 |
| | B27 | 1,365 | 0,703 | 2,157 | 1,022 | 0,558 | 1,402 |
| | BSA | Fixed to 1 (due to less experimental data in BSA condition) | | | | 0,948 | 2,301 |

| | | | | | | | |
|---|---|---|---|---|---|---|---|
| | HIMM-media | 0,572 | 1,981 | 1,013 | 0,943 | 0,741 | 1,317 |
| | ULT-media | 1,553 | 0,816 | 1,578 | 1,543 | 2,064 | 2,084 |

***List of donors and EndoC-βH5 batch condition alterations.***

*Step denotes how a condition affects a base parameter. The base parameter is affected through a multiplication, i.e. 1 means no change and 2 means double parameter value etc. The condition alteration ExenatidePotency affect both the negative and positive term in the potency equations (ExenatideInsulinDecay50 and ExenatideInsulin50). ExenatideResponse affect both the negative and positive term in the response magnitude equations (ExenatideInsulinDecayMax and ExenatideInsulinMax). Fislets0 affect the base parameter Fislets0. Note that Islet donors affect the base parameters describing the human islets, and the EndoC-βH5 batches affect the base parameters describing the EndoC-βH5 cultures. For merged cells, the same parameter is used to describe the media conditions.*

| Condition alteration | | **ExenatideResponse** | **ExenatidePotency** | **Filets0** |
|---|---|---|---|---|
| Description | | Exenatide max and min response amplitude | Exenatide $EC_{100}$ | Initial cell functionality |
| Unit | | step from base parameter (fold-increase) | | |
| Islet donor | **1** | 1,185 | 0,427 | 0,432 |
| | **2** | 1,233 | 1,059 | 1,005 |
| | **3** | 0,625 | 0,846 | 0,731 |
| | **4** | 0,815 | 1,607 | 1,208 |
| | **5** | 0,923 | 2,024 | 0,853 |
| | **6** | 1,072 | 1,252 | 0,857 |
| | **7** | 1,131 | 1,381 | 0,499 |
| | **8** | No data (fixed to 1) | | 0,793 |
| | **9** | No data (fixed to 1) | | 2,542 |
| | **10** | No data (fixed to 1) | | 1,938 |
| | **11 (validation Fig. 5C)** | No data (fixed to 1) | | 3,00 |
| EndoC-βH5 batch | **1 (spheroids)** | 0,652 | 1,086 | 0,670 |
| | **2 (spheroids)** | 0,523 | 0,694 | 1,488 |
| | **3 (spheroids)** | 3,036 | 1,271 | 0,909 |
| | **4 (spheroids)** | 0,457 | 0,772 | 0,849 |
| | **5 (monolayers)** | 1,345 | 0,731 | 1,010 |

***List of liver spheroid batch condition alterations.***

*The step indicates how a condition affects a base parameter. The base parameter is influenced through multiplication, where 1 means no change, 2 means doubling the parameter value, etc.The name of the condition alteration is identical to the base parameter which it affects, i.e. ISk in the table affects the base parameter ISk.*

| Condition alteration | | **Fspheroids50** | **ISk** | **Fspheroids0** |
|---|---|---|---|---|
| Description | | Cell function stability | Resistance to insulin sensitivity changes | Initial cell functionality |
| Unit | | step from base parameter (fold-increase) | | |
| Liver spheroid batches | **1** | 1,354 | 1,108 | 0,389 |
| | **2** | 0,847 | 1,111 | 0,451 |
| | **3** | 0,614 | 1,491 | 1,517 |
| | **4** | 1,172 | 0,632 | 1,262 |
| | **5** | 0,832 | 0,826 | 2,164 |
| | **6 (validation Fig. 5C)** | Fixed to 1 (because calibration was only done to first media change marked with 'x') | | 0,5 |

***List of uncertainty and calibration parameters.***
*θ denotes best agreement to data.*

| **Parameter** | **Unit** | **Description** | **θ** |
|---|---|---|---|
| ExperimentalInducedStress | h | Tissue transfer time for study 8 exenatide dose-response. This parameter describes the change from expected glucose dose-response (**Fig. 2E,** green) to a potential damaged state in the subsequent exenatide dose-response (**Fig. 2A,** green; low insulin secretion especially at the lower exenatide exposures) by decreasing the spheroid functionality equations. | 626,072 |
| InSpheroInsulinPhases_Uncertainty | fold-increase | Data extraction uncertainty for basal data point (**Fig. 2D**; 2.8 mM glucose) | 1,422 |
| FlowGSISBasalS | fold-increase | Allows for differences in basal insulin secretion when islets are in the external flow GSIS system | 4,5 |
| EndoCBasal_Uncertainty | fold-increase | Data extraction uncertainty for basal data point (Supplementary **Fig. S1D d,** 0 mM glucose) | 2,979 |
| WashingUncertaintyEndoC | fold-increase | Washing of EndoC-βH5 spheroids before static cell assays. A lower value means a bigger initial difference between intracellular and media concentration, i.e. more glucose is left after washing. The assumption was made that all washing was done with this uncertainty to minimise complexity and risk of overfitting. A value of 1 means no impact from the uncertainty parameter. | 0,981 |

#### c) Model ordinary differential equations (ODEs)

Four model states represent metabolites in media (**Eqs. S1–S4**).

$$\frac{\mathrm{d}}{\mathrm{dt}}(\mathrm{Glucose_{Medium}}) = Glucose_{IsletsToMedium}\frac{Vm_{Islets}}{Vm_{Medium}} + EGP\frac{Vm_{Hep}}{Vm_{Medium}} - (Uil + Glucose_{MediumToIslets}) \qquad \textbf{(Eq. S1)}$$

where $\mathrm{Glucose_{Medium}}$ describes glucose in the media, and where the components of the right-hand side are defined as:

$Glucose_{IsletsToMedium}$ denotes glucose transportation from pancreatic islets/spheroids to media,
$Glucose_{MediumToIslets}$ denotes glucose transportation from media to pancreatic islets/spheroids,
$EGP$ denotes glucose transportation from liver spheroids to media,
$Uil$ denotes glucose transportation from media to liver spheroids,
$Vm_{Islets}$ denotes pancreatic islet/spheroid volume,
$Vm_{Medium}$ denotes total media volume, and
$Vm_{Hep}$ denotes liver spheroid volume.

$$\frac{\mathrm{d}}{\mathrm{dt}}(\mathrm{Insulin_{Medium}}) = \mathrm{S}\frac{Vm_{Islets}}{Vm_{Medium}} - (InsulinClearance_{Liver}) \qquad \textbf{(Eq. S2)}$$

where $\mathrm{Insulin_{Medium}}$ describes insulin in the media, and where the components of the right-hand side are defined as:

$S$ denotes insulin secretion from the pancreatic islets/spheroids to media,
$InsulinClearance_{Liver}$ denotes liver spheroid insulin clearance,
$Vm_{Islets}$ denotes pancreatic islet/spheroid volume, and
$Vm_{Hep}$ denotes liver spheroid volume.

$$\frac{\mathrm{d}}{\mathrm{dt}}(\mathrm{Exenatide_{Medium}}) = -\,ExenatideDegradation \qquad \textbf{(Eq. S3)}$$

where $\mathrm{Exenatide_{Medium}}$ describes exenatide in the media, and where the components of the right-hand side are defined as:

$ExenatideDegradation$ denotes exenatide degradation from media.

$$\frac{\mathrm{d}}{\mathrm{dt}}(\mathrm{Lactate_{Medium}}) = 2 * LactateP\frac{Vm_{Hep}}{Vm_{Medium}} - (LactateU) \qquad \textbf{(Eq. S4)}$$

where $\mathrm{Lactate_{Medium}}$ describes lactate in the media, and where the components of the right-hand side are defined as:

$LactateP$ denotes glucose metabolised into lactate and transported from intracellular hepatic glucose compartment to media,
2 accounts for the mass balance between glucose (6 carbons) and lactate (3 carbons),
$LactateU$ denotes lactate metabolised from media into intracellular hepatic glucose,
$Vm_{Medium}$ denotes total media volume, and
$Vm_{Hep}$ denotes liver spheroid volume.

The hepatic metabolism is central to describe the observed lactate, insulin, and glucose concentrations in the media. To describe all three metabolites with a minimal model complexity, two key assumptions were made: a) the difference between intracellular and media lactate concentrations is negligible, and therefore lactate is secreted directly from the hepatic glucose compartment into the media lactate compartment; and b) the difference between intracellular and media insulin concentrations are negligible and therefore hepatic insulin clearance takes part in the ODE of $\mathrm{Insulin_{Medium}}$. Due to these two assumptions, intracellular hepatic metabolism can be described by one single ODE (**Eq. S5**).

$$\frac{\mathrm{d}}{\mathrm{dt}}\left(\mathrm{Glucose_{Spheroids}}\right) = \mathrm{Uil}\frac{Vm_{Medium}}{Vm_{Hep}} + 0.5 * LactateU\frac{Vm_{Medium}}{Vm_{Hep}} - (EGP + LactateP + Glucose_{out}) \qquad \textbf{(Eq. S5)}$$

where $\mathrm{Glucose_{Spheroids}}$ describes glucose in the liver spheroids, and where the components of the right-hand side are defined as:
Uil denotes glucose transportation from media into the liver spheroids,
$LactateU$ denotes lactate metabolised from media to the liver spheroids,
0.5 accounts for the mass balance between glucose (6 carbons) and lactate (3 carbons),
$EGP$ denotes glucose transportation out of liver spheroids to media,
$LactateP$ denotes glucose metabolised into lactate and transported from the liver spheroids to media,
$Glucose_{out}$ denotes glucose metabolised to non-lactate which is sent out of the model,
$Vm_{Medium}$ denotes total media volume, and
$Vm_{Hep}$ denotes liver spheroid volume.

In the model, insulin resistance can change depending on the intracellular hepatic glucose concentration. This glucose-dependent insulin resistance development is represented by an ODE that describes the diseased state of the liver spheroids. The diseased state increases when intracellular hepatic glucose concentrations are above a threshold (**Eq. S6**).

$$\frac{\mathrm{d}}{\mathrm{dt}}(\mathrm{GlucoseOverHealthy}) = \max\left(0, \mathrm{Glucose_{Spheroids}} - \mathrm{IR_{UpperLimit}}\right) \qquad \textbf{(Eq. S6)}$$

where GlucoseOverHealthy describes how much and for how long glucose has been greater than a specific threshold, and where the components of the right-hand side are defined as:
$\mathrm{Glucose_{Spheroids}}$ denotes intracellular hepatic glucose concentration, and
$\mathrm{IR_{UpperLimit}}$ denotes the threshold at which the diseased state GlucoseOverHealthy increases.

The ODEs that describe pancreatic metabolism are centralised around insulin production and secretion. In the model, insulin is first produced and stored in the pancreatic cells (**Eq. S7**), and the production of the insulin is dependent on pancreatic glucose concentrations (**Eq. S8**).

$$\frac{\mathrm{d}}{\mathrm{dt}}(\mathrm{Insulin_{Islets}}) = IP * \frac{1}{Vm_{Islets}} - (S) \qquad \textbf{(Eq. S7)}$$

where $\mathrm{Insulin_{Islets}}$ describes insulin in the pancreatic islets/spheroids, and where the components of the right-hand side are defined as:
$IP$ denotes insulin production,
$S$ denotes insulin secretion out of the islets/spheroids and into the media, and
$Vm_{Islets}$ denotes total pancreatic islets/spheroids volume.

$$\frac{\mathrm{d}}{\mathrm{dt}}(\mathrm{Glucose_{Islets}}) = Glucose_{MediumToIslets}\frac{Vm_{Medium}}{Vm_{Islets}} - (Glucose_{IsletsToMedium}) \qquad \textbf{(Eq. S8)}$$

where $\mathrm{Glucose_{Islets}}$ describes glucose in the pancreatic islets/spheroids, and where the components of the right-hand side are defined as:

$Glucose_{MediumToIslets}$ denotes glucose transportation from media to pancreatic islets/spheroids,
$Glucose_{IsletsToMedium}$ denotes glucose transportation from pancreatic islets/spheroids to media,
$Vm_{Islets}$ denotes pancreatic islet/spheroid volume, and
$Vm_{Medium}$ denotes total media volume.

In the calculation of insulin production, a delay is introduced between glucose rate of change and the glucose rate of change dependent insulin production (**Eq. S9**).

$$\frac{d}{dt}(\mathrm{IP_{ROCdelay}}) = \mathrm{IP_{ROCdelayK}} * (\mathrm{IP_{ROC}} - \mathrm{IP_{ROCdelay}}) \qquad \textbf{(Eq. S9)}$$

where $\mathrm{IP_{ROCdelay}}$ describes a delayed response between insulin production and glucose rate of change, and where the components of the right-hand side are defined as:
$\mathrm{IP_{ROCdelayK}}$ denotes the delay magnitude (where a higher value means a lower delay), and
$\mathrm{IP_{ROC}}$ denotes the glucose rate of change dependent insulin secretion before the delay.

In the model, pancreatic islet and spheroid proliferation is described using one ODE (**Eq. S10**).

$$\frac{d}{dt}(\mathrm{PancreticCells}) = \mathrm{GrowthRate} - \mathrm{DeathRate} \qquad \textbf{(Eq. S10)}$$

where PancreticCells describes the total amount of pancreatic cells in the study, and where the components of the right-hand side are defined as:
GrowthRate denotes pancreatic proliferation, and
DeathRate denotes pancreatic deathrate.

All ODEs listed above are defined by metabolic reactions which are one-by-one listed below.

#### d) Model reactions

Two metabolic reactions were used to describe total spheroid/islet volumes from total number of cells in experiment (**Eqs. S11-S12**).

$$\mathrm{Vm_{Hep}} = 3.4 * 10^{-11} * \mathrm{CountLiver} \qquad \textbf{(Eq. S11)}$$

where $\mathrm{Vm_{Hep}}$ describes total liver spheroid volume, and where the components of the right-hand side are defined as:
$3.4 * 10^{-11}$ denotes estimated single liver cell volume, and
CountLiver denotes estimated total liver cell number in experiment.

$$\mathrm{Vm_{Islets}} = 4.4 * 10^{-13} * \mathrm{PancreticCells} \qquad \textbf{(Eq. S12)}$$

where $\mathrm{Vm_{Islets}}$ describes total pancreatic islet/spheroid volume, and where the components of the right-hand side are defined as:
$4.4 * 10^{-13}$ denotes estimated single pancreatic cell volume, and
PancreticCells denotes total pancreas cell count in experiment.

The total cell volume equations (**Eqs. S11-S12**) therefore assumes that the volume scales one-to-one to total cell number, and that EndoC-βH5 cells and the human islet cells have the same volume. These two assumptions were considered to be sufficient for our applications, due to: a) the large difference in volume between the media and the spheroids/islets; and b) the un-rejected model agreement to data.

The total cell number was used to scale the full metabolic rates of the islets and spheroids between the studies (**Eqs. S13-S14**).

$$\text{AllometricScaling}_{\text{Hep}} = \mathit{LiverSpheroids}^{\mathit{AllometricLiverScalingK}} \qquad \textbf{(Eq. S13)}$$

where $\text{AllometricScaling}_{\text{Hep}}$ describes the total metabolic rate scaling, and where the components of the right-hand side are defined as:
$\mathit{LiverSpheroids}$ denotes liver spheroids in the experimental system, and
$\mathit{AllometricLiverScalingK}$ denotes the scaling constant.

$$\text{AllometricScaling}_{\text{Islets}} = \mathit{PancreaticIslets}^{\mathit{AllometricSPancreaticScalingK}} \qquad \textbf{(Eq. S14)}$$

where $\text{AllometricScaling}_{\text{Islets}}$ describes the total metabolic rates of pancreatic cultures, and where the components of the right-hand side are defined as:
$\mathit{PancreaticIslets}$ denotes pancreatic spheroids/islets in the experimental system, and
$\mathit{AllometricSPancreaticScalingK}$ denotes the scaling constant.

For the EndoC-βH5 monolayer simulations, the $\mathit{PancreaticIslets}$ (**Eq. S14**) was set to total cell number divided by 2000, where 2000 is the total number of cells assumed to be in our EndoC-βH5 spheroids. From the analysis of the EndoC-βH5 batch condition alterations (**Fig. 6B a**) we deem this assumption to be sufficient for our applications. This is based on condition alteration results, where batch ID 5 (which describes only the EndoC-βH5 monolayers) compared to batch ID 1-4 (which describes only the EndoC-βH5 spheroids) are not far from each other (**Fig. 6B a** yellow colour similar to the other colours). If the assumption would not be sufficient then the condition alteration would be far apart to compensate.

Insulin sensitivity is described with a static parameter, a glucose-dependent variable, and a time-dependent variable (**Eq. S15**).

$$IS = \text{IS0} + \text{ISk} * (\text{GlucoseOverHealthy} + \text{InsulinResistanceTimeDependency} * \text{Time}) \qquad \textbf{(Eq. S15)}$$

where $IS$ describes a multiplication of hepatic insulin response ($EC_{50}$), and where the components of the right-hand side are defined as:
IS0 denotes an initial hepatic insulin response,
ISk denotes volatility of insulin resistance development,
GlucoseOverHealthy denotes the disease progression state,
InsulinResistanceTimeDependency denotes insulin resistance progression dependency on time, and
Time denotes hours since experiment start.
Note that IS0 and InsulinResistanceTimeDependency are affected by media condition alterations, and that ISk is affected by liver spheroid batch condition alterations.

As the cells are studied *in vitro*, the metabolic rates of the islets and spheroids may decrease over time. This decrease in metabolic rates is described as functionality loss decay throughout this analysis. The functionality decay is assumed to depend on time. The functionality relation to time is affected by media condition alterations, which means that media conditions are allowed to affect functional baseline (Fislets0), how long before the functionality decay starts ($Fislets50$), and how rapid the decay is ($islets_{Hill}$). Furthermore, the initial functional baseline is also affected by batch and donor condition alterations (**Eqs. S16-S17**).

$$F_{Islets} = \text{Fislets0} - \frac{Fislets0 * Time^{Fislets_{Hill}}}{Fislets50^{Fislets_{Hill}} + Time^{Fislets_{Hill}}} \quad \textbf{(Eq. S16)}$$

where $F_{Islets}$ describes loss of pancreatic islets/spheroids metabolic functionality, and where the components of the right-hand side are defined as:
Fislets0 denotes initial functionality,
$Time$ denotes hours since experiment start,
$Fislets_{Hill}$ denotes hill coefficient of functionality decay rate, and
$Fislets50$ denotes pancreas islets/spheroids stability. Note that the parameters Fislets0, $Fislets50$, and $Fislets_{Hill}$ are affected by media condition alterations, and that the parameter Fislets0 is affected by donor and batch condition alterations.

$$F_{Spheroids} = \text{Fspheroids0} - \frac{Fspheroids0 * Time^{Fspheroids_{Hill}}}{Fspheroids50^{Fspheroids_{Hill}} + Time^{Fspheroids_{Hill}}} \quad \textbf{(Eq. S17)}$$

where $F_{Spheroids}$ describes changes to liver spheroid functionality, and where the components of the right-hand side are defined as: $Fspheroids0$ denotes initial functionality,
$Time$ denotes hours since experiment start,
$Fspheroids_{Hill}$ denotes functionality decay rate, and
$Fspheroids50$ denotes pancreas islets/spheroids stability until loss of functionality. Note that the parameters Fspheroids0, $Fspheroids50$, and $Fspheroids_{Hill}$ are affected by media condition alterations, and that the parameter Fspheroids0 is affected by batch condition alterations.

Hepatic insulin response is described with a dependency on insulin (**Eq. S18**).

$$InsulinResponse = \frac{\text{InsulinResponseMax} * Insulin_{Medium}^{InsulinResponse_{Hill}}}{(IS * InsulinResponse50)^{InsulinResponse_{Hill}} + Insulin_{Medium}^{InsulinResponse_{Hill}}} \quad \textbf{(Eq. S18)}$$

where $InsulinResponse$ describes an increase in insulin dependent reaction rates, and where the components of the right-hand side are defined as:
InsulinResponseMax denotes maximum response amplitude,
$Insulin_{Medium}$ denotes insulin concentration in media,
$InsulinResponse_{Hill}$ denotes hill coefficient of the change from no response to maximum response,
$InsulinResponse50$ denotes insulin exposure when $InsulinResponse$ reaches half of maximum effect, and
$IS$ denotes insulin resistance progression in form of a factor greater or equal to 1 that reduces the potency of the insulin response.

The hepatic insulin response increases glucose utilisation, which is separated into lactate production (**Eq. S19**), and non-lactate production (**Eq. S20**).

$InsulinResponse_{lactate} = 1 + InsulinResponse * InsulinResponseLactateK$ (**Eq. S19**)
where $InsulinResponse_{lactate}$ describes insulin response of lactate production, and where the components of the right-hand side are defined as:
$InsulinResponse$ denotes the general hepatic insulin response,
1 represent the baseline level in the absence of insulin, and
$InsulinResponseLactateK$ denotes the fraction of the $InsulinResponse$ which affects the glucose to lactate reaction rate.

$InsulinResponse_{Gout} = 1 + InsulinResponse * (1 - InsulinResponseLactateK)$ (**Eq. S20**)
where $InsulinResponse_{Gout}$ describes hepatic glucose utilisation not metabolised into lactate, and where the components of the right-hand side are defined as:
$InsulinResponse$ denotes the general hepatic insulin response,
1 represent the baseline level in the absence of insulin, and
$InsulinResponseLactateK$ denotes the fraction of the $InsulinResponse$ affecting glucose metabolised into lactate.

Exenatide loss was described with a constant half-life (**Eq. S21**).

$\mathrm{Exenatide_{Degradation}} = \mathrm{Exenatide_{DegradationK}} * \mathrm{Exenatide_{Medium}}$ (**Eq. S21**)
where $\mathrm{Exenatide_{Degradation}}$ describes exenatide half-life, and where the components of the right-hand side are defined as:
$\mathrm{Exenatide_{DegradationK}}$ denotes loss of exenatide over time and
$\mathrm{Exenatide_{Medium}}$ denotes concentration of exenatide in media.

The effect of exenatide on insulin secretion and production was described with a dependency on exenatide in the media (**Eq. S22**). The effect increases insulin secretion (**Eq. S23**), glucose rate of change dependent insulin production (**Eq. S24**), and glucose concentration dependent insulin production (**Eq. S25**).

$$\mathrm{Exenatide_{PD}} = \frac{ExenatideInsulinMax * Exenatide_{Medium}^{Exenatide_{Hill}}}{ExenatideInsulin50^{Exenatide_{Hill}} + Exenatide_{Medium}^{Exenatide_{Hill}}} - \frac{ExenatideInsulinDecayMax * Exenatide_{Medium}^{Exenatide_{Hill}}}{ExenatideInsulinDecay50^{Exenatide_{Hill}} + Exenatide_{Medium}^{Exenatide_{Hill}}} \quad \textbf{(Eq. S22)}$$

where $\mathrm{Exenatide_{PD}}$ describes the bell-shaped exenatide dose-response effect on insulin secretion and production, and where the components of the right-hand side are defined as:
$ExenatideInsulinMax$ and $ExenatideInsulinDecayMax$ denote the maximum effect of both the positive and negative parts of the bell-shaped dose-response curve,
$Exenatide_{Hill}$ denotes hill coefficient from no response to maximum response,
$Exenatide_{Medium}$ denotes exenatide concentration in the media,
$ExenatideInsulin50$ and $ExenatideInsulinDecay50$ denote $EC_{50}$ and $IC_{50}$ of the bell-shaped dose-response curve. Note that $ExenatideInsulin50$, $ExenatideInsulinDecay50$, $ExenatideInsulinMax$, and $ExenatideInsulinDecayMax$ are affected by media, batch, and donor condition alterations.

$$\text{Exenatide}_{\text{S}} = 1 + \text{Exenatide}_{\text{PD}} \qquad \textbf{(Eq. S23)}$$

Where $\text{Exenatide}_{\text{S}}$ describes exenatide effect on insulin secretion, and where the components of the right-hand side are defined as:
1 represent the baseline level in the absence of exenatide,
$\text{Exenatide}_{\text{PD}}$ denotes the overall exenatide effect.

$$\text{Exenatide}_{\text{IProc}} = 1 + \text{Exenatide}_{\text{PD}} * (1 - \text{Exenatide}_{\text{IP}}) \qquad \textbf{(Eq. S24)}$$

where $\text{Exenatide}_{\text{IProc}}$ describes exenatide effect on phase-I insulin production, and where the components of the right-hand side are defined as:
1 represent the baseline level in the absence of exenatide,
$\text{Exenatide}_{\text{PD}}$ denotes general exenatide effect, and
$\text{Exenatide}_{\text{IP}}$ denotes ratio of exenatide effect which affects the phase-I/phase-II insulin production.

$$\text{Exenatide}_{\text{IPbasal}} = 1 + \text{Exenatide}_{\text{PD}} * \text{Exenatide}_{IP} \qquad \textbf{(Eq. S25)}$$

where $\text{Exenatide}_{\text{IPbasal}}$ describes exenatide effect on phase-II insulin production, and where the components of the right-hand side are defined as:
1 represent the baseline level in the absence of exenatide,
$\text{Exenatide}_{\text{PD}}$ denotes general exenatide effect, and
$\text{Exenatide}_{\text{IP}}$ denotes ratio of exenatide effect which affects the phase-I/phase-II insulin production.

The mathematical description of pancreas-liver biology includes pancreatic proliferation. In the data-driven hypothesis testing an exenatide impact on this proliferation was tested with a dependency on exenatide concentration in media (**Eq. S26**).

$$\text{Exenatide}_{\text{Stabilisation}} = 1 + \frac{\text{ExenatideStabilisationMax} * Exenatide_{Medium}}{\text{ExenatideStabilisationEC50} + Exenatide_{Medium}} \qquad \textbf{(Eq. S26)}$$

where $\text{Exenatide}_{\text{Stabilisation}}$ describes exenatide effect on pancreatic growth rate, and where the components of the right-hand side are defined as:
1 represent the baseline level in the absence of exenatide,
ExenatideStabilisationMax denotes maximum effect on growth rate,
$Exenatide_{Medium}$ denotes exenatide concentration in media, and
ExenatideStabilisationEC50 denotes when effect reaches half of the maximum stabilisation. This effect on growth rate could not be concluded from the data (**Supplementary Fig. S1A b,** right).

Glucose transportation between media and liver spheroids is described with a dependency on glucose in the media and in the spheroids (**Eqs. S27-S28**).

$$Uil = \text{F}_{\text{Spheroids}} * \text{AllometricLiverScaling} * \text{Uil}_{\text{iiK}} * \frac{\text{Glucose}_{\text{Medium}}}{Vm_{Medium}} \qquad \textbf{(Eq. S27)}$$

where $Uil$ describes glucose transportation from the media to liver spheroids, and where the components of the right-hand side are defined as:
$\text{F}_{\text{Spheroids}}$ denotes liver spheroid functionality,
AllometricLiverScaling denotes metabolic rate scaling to total cell number,
$\text{Uil}_{\text{iiK}}$ denotes transportation rate,

$\text{Glucose}_{\text{Medium}}$ denotes glucose in media, and
$Vm_{Medium}$ denotes volume in media to take into consideration volume differences between the compartments.

$$EGP = * \text{Uil}_{\text{iiK}} * \text{Glucose}_{\text{Spheroids}} * \frac{\text{F}_{\text{Spheroids}} * \text{AllometricLiverScaling}}{Vm_{Hep}} \quad \textbf{(Eq. S28)}$$

where $EGP$ describes glucose transportation from liver spheroids to media, and where the components of the right-hand side are defined as: $\text{F}_{\text{Spheroids}}$ denotes liver spheroid functionality,
AllometricLiverScaling denotes metabolic rate scaling to total cell number,
$\text{Uil}_{\text{iiK}}$ denotes transportation rate,
$\text{Glucose}_{\text{Spheroids}}$ denotes glucose in liver spheroids, and
$Vm_{Hep}$ denotes volume in spheroids to take into consideration volume differences between the compartments.

Hepatic lactate production is described with a dependency on insulin and hepatic glucose levels (**Eq. S29**) and hepatic lactate utilisation was described to be insulin independent (**Eq. S30**).

$$LactateProduction = \text{InsulinResponseLactate} * \text{LactateProductionK} * \text{Glucose}_{\text{Spheroids}} * \frac{\text{F}_{\text{Spheroids}} * \text{AllometricLiverScaling}}{Vm_{Hep}} \quad \textbf{(Eq. S29)}$$

where $LactateProduction$ describes lactate production from liver spheroids to media, and where the components of the right-hand side are defined as:
$\text{F}_{\text{Spheroids}}$ denotes liver spheroid functionality,
AllometricLiverScaling denotes metabolic rate scaling to total cell number,
InsulinResponseLactate denotes insulin response in the lactate production,
LactateProductionK denotes insulin independent reaction rate,
$\text{Glucose}_{\text{Spheroids}}$ denotes glucose in liver spheroids, and
$Vm_{Hep}$ denotes volume in spheroids to take into consideration volume differences between the compartments.

$$LactateU = \text{LactateUk} * \text{Lactate}_{\text{Medium}} * \frac{\text{F}_{\text{Spheroids}} * \text{AllometricLiverScaling}}{Vm_{Medium}} \quad \textbf{(Eq. S30)}$$

where $LactateU$ describes lactate utilisation of the hepatic cells from media, and where the components of the right-hand side are defined as:
$\text{F}_{\text{Spheroids}}$ denotes liver spheroid functionality,
AllometricLiverScaling denotes metabolic rate scaling to total cell number,
LactateUk denotes metabolic rate,
$\text{Lactate}_{\text{Medium}}$ denotes lactate in media, and
$Vm_{Medium}$ denotes media volume to take into consideration volume differences between the compartments.

Hepatic glucose utilisation into non-lactate was described with a dependency on insulin and hepatic glucose levels (**Eq. S31**).

$$Gout = \text{InsulinResponseGout} * \text{GoutK} * \text{Glucose}_{\text{Spheroids}} * \frac{\text{F}_{\text{Spheroids}} * \text{AllometricLiverScaling}}{Vm_{Hep}}$$

**(Eq. S31)**

where $Gout$ describes glucose utilised into metabolites that are not lactate, and where the components of the right-hand side are defined as: $\text{F}_{\text{Spheroids}}$ denotes liver spheroid functionality,
AllometricLiverScaling denotes metabolic rate scaling to total cell number,
InsulinResponseGout denotes insulin response of non-lactate metabolites,
GoutK denotes insulin independent reaction rate,
$\text{Glucose}_{\text{Spheroids}}$ denotes glucose in liver spheroids, and
$Vm_{Hep}$ denotes volume in spheroids to take into consideration volume differences between the compartments.

Hepatic insulin clearance was described with a dependency on insulin concentrations (**Eq. S32**).

$$InsulinClearanceLiver = \text{InsulinClearanceLiverk} * \text{Insulin}_{\text{Medium}} * \frac{\text{F}_{\text{Spheroids}} * \text{AllometricLiverScaling}}{Vm_{Medium}}$$

**(Eq. S32)**

where $InsulinClearanceLiver$ describes hepatic insulin clearance, and where the components of the right-hand side are defined as: $\text{F}_{\text{Spheroids}}$ denotes liver spheroid functionality,
AllometricLiverScaling denotes metabolic rate scaling to total cell number, InsulinClearanceLiverk denotes insulin clearance metabolic rate,
$\text{Insulin}_{\text{Medium}}$ denotes insulin in media, and
$Vm_{Medium}$ denotes media volume to take into consideration volume differences between the compartments.

Glucose transportation between pancreatic islets/spheroids and media is described to be dependent on glucose concentrations in the islets/spheroids and media (**Eqs. S33-S34**).

$$GlucoseMediumToIslets = \text{GlucoseIsletsMediumDiffusion} * \text{Glucose}_{\text{Medium}} * \frac{\text{F}_{\text{islets}} * \text{AllometricPancreasScaling}}{Vm_{Medium}}$$

**(Eq. S33)**

where $GlucoseMediumToIslets$ describes glucose transportation from media to pancreatic islets/spheroids, and where the components of the right-hand side are defined as:
$\text{F}_{\text{islets}}$ denotes pancreatic islet/spheroid spheroid functionality,
AllometricPancreasScaling denotes metabolic rate scaling to total cell number,
GlucoseIsletsMediumDiffusion denotes glucose transportation rate,
$\text{Glucose}_{\text{Medium}}$ denotes glucose in media, and
$Vm_{Medium}$ denotes media volume to take into consideration volume differences between the compartments.

$$GlucoseIsletsToMedium = \text{GlucoseIsletsMediumDiffusion} * \text{Glucose}_{\text{Islets}} * \frac{\text{F}_{\text{Islets}} * \text{AllometricPancreasScaling}}{Vm_{Islets}}$$

**(Eq. S34)**

where $GlucoseIsletsToMedium$ describes glucose transportation from pancreatic islets/spheroids to media, and where the components of the right-hand side are defined as:
$\text{F}_{\text{islets}}$ denotes pancreatic islet/spheroid functionality,

AllometricPancreasScaling denotes metabolic rate scaling to total cell number,
GlucoseIsletsMediumDiffusion denotes transportation rate,
$\mathrm{Glucose}_{\mathrm{Islets}}$ denotes glucose in islets/spheroids, and
$Vm_{Islets}$ denotes pancreatic islet/spheroid volume to take into consideration volume differences between the compartments.

Pancreatic insulin production is described with a dependency on pancreatic glucose rate of change (**Eqs. S35-S36**), dependency on pancreatic glucose levels (**Eq. S37**) and with a basal insulin production (**Eq. S38**) as a sum (**Eq. S39**).

$$GlucoseROC = \max(0, \frac{\mathrm{d}}{\mathrm{dt}}(\mathrm{Glucose}_{\mathrm{Islets}})) \quad \textbf{(Eq. S35)}$$

where $GlucoseROC$ describes pancreatic glucose rate of change, and where the components of the right-hand side are defined as:
max denotes that the function is always positive or zero, and
$\frac{\mathrm{d}}{\mathrm{dt}}(\mathrm{Glucose}_{\mathrm{Islets}})$ denotes intracellular pancreatic glucose rate of change.

$$IP_{ROC} = \mathrm{F}_{\mathrm{islets}} * \mathrm{AllometricPancreasScaling} * (\mathrm{IP}_{\mathrm{ROCk}} * \mathrm{GlucoseROC}) * Exenatide_{IProc} \quad \textbf{(Eq. S36)}$$

where $IP_{ROC}$ describes glucose rate of change dependent insulin production, and where the components of the right-hand side are defined as:
$\mathrm{F}_{\mathrm{islets}}$ denotes pancreatic islet/spheroid functionality,
AllometricPancreasScaling denotes metabolic rate scaling to total cell number,
$\mathrm{IP}_{\mathrm{ROCk}}$ denotes insulin production scaling of the positive rate of change,
GlucoseROC denotes positive pancreatic glucose rate of change, and
$Exenatide_{IProc}$ denotes exenatide effect on glucose rate of change dependent insulin production.

$$IP_{Basline} = \max(0, \mathrm{Glucose}_{\mathrm{Islets}} - \mathrm{IP}_{\mathrm{BaslineK}})$$

where $IP_{Basline}$ describes pancreatic glucose concentration is above a threshold, and where the components of the right-hand side are defined as:
max denotes that the function is always positive or zero,
$\mathrm{Glucose}_{\mathrm{Islets}}$ denotes glucose in the islets/spheroids, and
$\mathrm{IP}_{\mathrm{BaslineK}}$ denotes the threshold glucose needs to be above to start triggering the glucose concentration dependent insulin production.

$$\mathrm{IP}_{\mathrm{Baseline}} = \mathrm{F}_{\mathrm{Islets}} * \mathrm{AllometricPancreaseScaling} * \frac{IP_{BaselineMax} * IP_{Basline}{}^{IpBaseline_{Hill}}}{IPBaseline50^{IpBaseline_{Hill}} + IP_{Basline}{}^{IpBaseline_{Hill}}} * \mathrm{Exenatide}_{\mathrm{IPbasal}} \quad \textbf{(Eq. S37)}$$

where $\mathrm{IP}_{\mathrm{Baseline}}$ describes glucose concentration dependent insulin production, and where the components of the right-hand side are defined as:
$\mathrm{F}_{\mathrm{islets}}$ denotes pancreatic islet/spheroid functionality,
AllometricPancreasScaling denotes metabolic rate scaling to total cell number,
$IP_{BaselineMax}$ denotes maximum glucose concentration dependent insulin production,
$IpBaseline_{Hill}$ denotes hill coefficient between no glucose concentration dependent insulin production to maximum,

$IPBaseline50$ denotes at what glucose exposure half of the maximum production is, and $Exenatide_{IProc}$ denotes exenatide effect on glucose concentration dependent insulin production.

$\mathrm{IP_{Basal}} = \mathrm{F_{Islets}} * \mathrm{AllometricPancreaseScaling} * \mathrm{IP_{basalK}}$ **(Eq. S38)**
where $\mathrm{IP_{Basal}}$ describes basal insulin production, and where the components of the right-hand side are defined as:
$\mathrm{F_{islets}}$ denotes pancreatic islet/spheroid functionality,
AllometricPancreasScaling denotes metabolic rate scaling to total cell number, and
$\mathrm{IP_{basalK}}$ denotes the basal insulin production.

$\mathrm{IP} = \mathrm{IP_{basal}} + IP_{ROCdelay} + IP_{baseline}$ **(Eq. S39)**
where IP describes the full insulin production, which is a sum of:
$\mathrm{IP_{basal}}$ denotes the basal insulin production,
$IP_{ROCdelay}$ denotes the delayed glucose rate of change dependent insulin production, and
$IP_{baseline}$ denotes the glucose concentration dependent insulin production.

Pancreatic insulin secretion is described with a dependency on stored insulin and exenatide effect (**Eq. S40**).

$S = \frac{\mathrm{S_{max}} * Insulin_{Islets}}{\mathrm{S50} + Insulin_{Islets}} * \mathrm{Exenatide_S} * \frac{\mathrm{F_{Islets}} * \mathrm{AllometricPancreaseScaling}}{Vm_{Islets}}$ **(Eq. S40)**
where S describes insulin secretion from the pancreatic cells to the media, and where the components of the right-hand side are defined as:
$\mathrm{F_{islets}}$ denotes pancreatic islet/spheroid functionality,
AllometricPancreasScaling denotes metabolic rate scaling to total cell number,
$\mathrm{S_{max}}$ denotes maximum secretion,
$Insulin_{Islets}$ denotes pancreatic insulin concentration,
S50 denotes at what insulin exposure half of the maximum secretion is reached,
$\mathrm{Exenatide_S}$ denotes exenatide effect on insulin secretion, and
$Vm_{Islets}$ denotes islet/spheroid volume to take into consideration volume differences between the compartments.

Pancreatic proliferation is described with a dependency on glucose and exenatide as the difference between growth rate (**Eq. S41**) and death rate (**Eq. S42**).

$GrowthRate = kv * r1 * Glucose_{Islets} * Exenatide_{Stabilisaiton}\ \mathrm{F_{Islets}} * \mathrm{AllometricPancreaseScaling}$ **(Eq. S41)**
where $GrowthRate$ denotes pancreatic proliferation, and where the components of the right-hand side are defined as:
$\mathrm{F_{islets}}$ denotes pancreatic islet/spheroid functionality,
AllometricPancreasScaling denotes metabolic rate scaling to total cell number,
$kv$ denotes proliferation volatility,
$r1$ denotes positive growth rate dependency on pancreatic glucose concentration,
$Glucose_{Islets}$ denotes the islet/spheroid glucose concentration, and
$Exenatide_{Stabilisaiton}$ denotes exenatide effect on growth rate.

$DeathRate = kv * (d0 + (r2 * Glucose_{Islets})^2) * \mathrm{F_{Islets}} * \mathrm{AllometricPancreaseScaling}$ **(Eq. S42)**
where $DeathRate$denotes describes pancreatic cell death, and where the components of the right-hand side are defined as:
$\mathrm{F_{islets}}$ denotes pancreatic islet/spheroid functionality,
$\mathrm{AllometricPancreasScaling}$ denotes metabolic rate scaling to total cell number,
$kv$ denotes proliferation volatility,
$d0$ denotes basal death rate,
$r2$ denotes death rate dependency on pancreatic glucose concentration $Glucose_{Islets}$.
