## Supplementary Note S2 for "An Integrated Analysis of GLP-1R Agonist Mechanisms: Addressing Study Variations in Heterogeneous Cell Systems"

This supplementary note provides additional reasoning for two studies in which no exenatide response was observed.

### Discussion

A lack of exenatide effect was observed in two studies (human islets: Study 15, **Supplementary Fig. S2A**; EndoC-βH5 spheroids: Study 16, **Supplementary Fig. S2B**). Data from both studies are provided in the supplementary material (**Supplementary Tables S17–S18**). Several potential explanations for the absence of an exenatide effect were hypothesized. Four possible hypotheses were considered:

1. **Batch-to-batch and donor-to-donor variability**: It was initially hypothesized that variability between batches or donors could explain the lack of response. However, this was ruled out for the human islets (lot: hIsMT 196), as they had been pre-characterized for GLP-1 receptor responsiveness prior to purchase. This hypothesis was also rejected for EndoC-βH5 spheroids, since the EndoC-βH5 cell lot (202201–220614; production date: 2022-07-05) and spheroid batch (batch 3) had demonstrated a clear exenatide effect in another study (Study 8; **Fig. 2A**, green).
2. **Hyperglycemia toxicity**: Another hypothesis was that exenatide responsiveness might be lost due to hyperglycaemic toxicity. Discussions with human islet suppliers indicated that such effects have been observed previously. We found this explanation unlikely in our case, as the human islet responsiveness to exenatide was assessed in static cell assays prior to exposure of hyperglycaemia (**Supplementary Fig. S2A a**). In addition, exenatide effects were observed following hyperglycaemic exposure in other experiments, including Study 4 (human islets) and Study 11 (EndoC-βH5 spheroids). The mathematical modelling provided sufficient agreement with all human cell data without incorporating a hyperglycaemia-induced reduction in exenatide responsiveness, indicating that such an interaction cannot be concluded from the data.
3. **FCS supplementation**: It was also hypothesized that fetal calf serum (FCS) supplementation could reduce the exenatide response. This hypothesis was not supported, as exenatide effects in the presence of FCS were observed in both human islet studies (e.g., Study 4) and EndoC-βH5 spheroid studies (e.g., Study 7). Furthermore, our mathematical modelling indicated that FCS enhanced exenatide efficacy compared to B27 supplementation (**Fig. 4A,** middle), indicating that an inhibitory effect of FCS on exenatide responsiveness cannot be concluded from the data.
4. **External factors**: Finally, it was hypothesized that external factors, such as contamination or damage to the islets or spheroids, could explain the lack of exenatide response. After ruling out the other three hypotheses, external factors were considered the most likely explanation. As these factors could not be incorporated into the modelling due to insufficient information regarding their nature, the two studies without an observed exenatide effect were excluded from mathematical modelling.
